# Multi-omics integration of scRNA-seq time series data predicts new intervention points for Parkinson’s disease

**DOI:** 10.1101/2023.12.12.570554

**Authors:** Katarina Mihajlović, Gaia Ceddia, Noël Malod-Dognin, Gabriela Novak, Dimitrios Kyriakis, Alexander Skupin, Nataša Pržulj

**Affiliations:** Barcelona Supercomputing Center (BSC), 08034 Barcelona, Spain; Department of Computer Science, University College London, WC1E 6BT London, United Kingdom; The Integrative Cell Signalling Group, Luxembourg Centre for Systems Biomedicine (LCSB), University of Luxembourg, Esch-sur-Alzette, Luxembourg; Luxembourg Institute of Health (LIH), Esch-sur-Alzette, Luxembourg; University of California San Diego, La Jolla, CA 92093, USA; ICREA, Pg. Lluís Companys 23, 08010 Barcelona, Spain

## Abstract

Parkinson’s disease (PD) is a complex neurodegenerative disorder without a cure. The onset of PD symptoms corresponds to 50% loss of midbrain dopaminergic (mDA) neurons, limiting early-stage understanding of PD. To shed light on early PD development, we study time series scRNA-seq datasets of mDA neurons obtained from patient-derived induced pluripotent stem cell differentiation. We develop a new data integration method based on Non-negative Matrix Tri-Factorization that integrates these datasets with molecular interaction networks, producing condition-specific “gene embeddings”. By mining these embeddings, we predict 193 PD-related genes that are largely supported (49.7%) in the literature and are specific to the investigated *PINK1* mutation. Enrichment analysis in Kyoto Encyclopedia of Genes and Genomes pathways highlights 10 PD-related molecular mechanisms perturbed during early PD development. Finally, investigating the top 20 prioritized genes reveals 12 previously unrecognized genes associated with PD that represent interesting drug targets.

## 1 Introduction

Parkinson’s disease (PD) is a complex multifactorial disease and the second most prevalent neurodegenerative disorder affecting about 2-3% of the population over the age of 65 (Poewe *et al*., 2017). Due to an ageing society, PD will continue to increase its burden on social systems and economy. In the United States alone, it is projected that by 2037 PD will impact more than 1.6 million individuals, surpassing the economic burden of $79 billion (Yang *et al*., 2020). PD’s exact cause is unknown, with only 5-10% of cases being linked to identified genetic mutations, while the remainder are categorized as idiopathic, lacking a known cause (Balestrino and Schapira, 2020). The current intervention strategies are limited to treating symptoms, and no curative treatment is available (Balestrino and Schapira, 2020). PD is characterized by the intracellular accumulation of misfolded *α*-synuclein proteins called Lewy bodies and the subsequent death of midbrain dopaminergic (mDA) neurons in the substantia nigra part of the brain (Balestrino and Schapira, 2020). Furthermore, increasing evidence implicates multiple molecular mechanisms in the disease, including disrupted mitochondrial function, calcium and protein homeostasis as well as oxidative and endoplasmic reticulum stress (Balestrino and Schapira, 2020; Poewe *et al*., 2017).

One of the main challenges in studying PD is the availability of tissue samples, as 60% of the mDA neurons have already died by the time of the diagnosis and 90% at the later stages of the disease (Giguère *et al*., 2018). This issue is limiting our understanding of the early stage of PD development. The recent technology of cellular reprogramming provides an alternative way of obtaining mDA neurons by converting the somatic cells of PD patients carrying disease-associated mutations into induced pluripotent stem cells (iPSCs) and differentiating them into mDA neurons (Lee *et al*., 2020; Smajić *et al*., 2022; Novak *et al*., 2022). This technique provides a practically unlimited source of mDA neurons that can be studied to uncover the molecular mechanisms driving PD. Recently, we used mDA neurons differentiated from iPSCs in the early stages of neural development (i.e., before PD is established) to investigate PD mechanisms by applying single-cell (SC) RNA sequencing (scRNA-seq) (Novak *et al*., 2022).

The emergence of SC sequencing techniques has led to an explosion of high-throughput measurements that can investigate cellular heterogeneity, offering the opportunity to study individual mDA neurons primed for degeneration. Different SC sequencing techniques have been applied to study the cellular response of heterogeneous cell types and further understand the molecular mechanisms underlying PD pathology and other diseases (Lee *et al*., 2020; Smajić *et al*., 2022; Novak *et al*., 2022). However, SC measurements are associated with high levels of noise that impede the subsequent data analysis. This challenge is often addressed with data integration strategies combining multi-omics data sets (as seen in the review of Luecken *et al*. (2022)). Such integrated data is often subjected to cell-level downstream analysis, including cell clustering, identifying cell types, and trajectory inference (Luecken and Theis, 2019). On the other hand, gene-level analysis (e.g., identifying disease gene markers) is typically based on identifying differentially expressed genes (DEGs) (Novak *et al*., 2022; Luecken and Theis, 2019; Welch *et al*., 2019). However, DEG analyses cannot uncover, for example, disease-related genes whose expressed proteins do not have altered expression but have undergone post-translational modifications, leading to disease pathogenesis (Thygesen *et al*., 2018). Furthermore, these approaches are also not adapted to properly handle time-series SC data as they uncover DEGs of each time point individually and do not provide a well-defined framework for predicting genes from all time points collectively. This prompts the need to develop novel non-DEG-based integration methods to discover new disease markers by analyzing SC time-series data. To jointly analyze time-series scRNA-seq data and identify different cell types, Jung *et al*. (2020) suggested a Non-Negative Matrix Factorization (NMF)-based approach.

Matrix factorization techniques (such as NMF and its extension Non-Negative Matrix Tri-Factorization (NMTF)) are popular co-clustering, dimensionality reduction and inference methods recently gaining attention for data integration. They project the original highly-dimensional data into lower-dimensional embedding spaces that are easier to handle and analyze (Yang *et al*., 2008). NMF and NMTF have been widely used for bulk data analysis to study, for instance, molecular networks (i.e. networks that capture relevant information about cellular functions and pathways) to suggest novel cancer-related genes (Malod-Dognin *et al*., 2019), protein functions (Peng *et al*., 2019) and drug-repurposing options (Tang *et al*., 2021). NMF-based approaches have also shown promising results in dealing with sparse SC samples (Welch *et al*., 2019; Argelaguet *et al*., 2020; Jung *et al*., 2020; Huizing *et al*., 2023). Furthermore, NMF-based methods have been used to jointly integrate SC data with molecular networks to identify types of SCs (Elyanow *et al*., 2020), to discover interpretable gene programs (Kunes *et al*., 2023) and to generate protein representations within various cellular contexts to identify therapeutic targets and nominate cell type contexts for rheumatoid arthritis and inflammatory bowel diseases (Li *et al*., 2023). Using matrix factorization approaches to integrate SC data with molecular networks (i.e., prior knowledge) allows us to benefit from the biologically relevant information in molecular networks and simultaneously minimize the inherent noisiness of SC data. Despite the advances in SC data analysis, no existing method is designed to uncover novel disease gene markers while fully exploiting time-series SC data and the information in prior knowledge contained in molecular interaction networks.

Here, we propose a new NMTF-based method, NetSC-NMTF, which simultaneously decomposes a time point-specific scRNA-seq dataset of a cell line harbouring a PD-associated mutation in the *PINK1* gene (I368N mutation), or a control one, with prior knowledge – protein-protein interaction (PPI), gene co-expression (COEX), metabolic interaction (MI), and genetic interaction (GI) networks. NetSC-NMTF produces gene embedding vectors (i.e., “gene embeddings”) that are biologically relevant, as shown by clustering and enrichment analysis in biological annotations from Gene Ontology (GO) (Ashburner *et al*., 2000), KEGG pathways (KP) (Kanehisa *et al*., 2017) and Reactome pathways (RP) (Jassal *et al*., 2020). Then, we introduce a 2-step downstream method that mines the “gene embeddings” across all cell conditions, identifying 193 PD-related gene predictions, of which 49.7% are associated with PD in the literature. Furthermore, in contrast to previous studies on SC data, our workflow reveals PD-associated genes beyond the standard DEG analysis (Novak *et al*., 2022; Welch *et al*., 2019). As the literature indicates that PD is a metabolic disease (Anandhan *et al*., 2017), we relate our 193 gene predictions to metabolic pathways by performing an enrichment analysis in KPs (Kanehisa *et al*., 2017). We highlight 10 significantly enriched KPs whose impairments in PD are supported by the literature, shedding light on the metabolic mechanisms that drive the progression of PD. Then, we manually validate the top 20 highest-scoring predictions to propose 12 new and promising PD-associated genes that include seven known and two potential new drug targets, representing potential candidates for developing novel treatments for PD. Finally, we demonstrate that the predictions are not only associated with PD, but are specific to the *PINK1* mutation. The methodological pipeline presented here is a flexible framework that could be extended to incorporate other types of SC, or bulk data and applied to other complex diseases.

## 2 Materials and Methods

### 2.1 Datasets

#### 2.1.1 Expression matrices and molecular networks

From Novak *et al*. (2022), we obtain the SC dataset that contains normalized scRNA-seq data of mDA neurons of two cell lines: a Parkinson’s disease cell line obtained from a 64-year-old male with a homozygous ILE368ASN mutation (P.I368N/P.I368N) in the *PINK1* gene and an age- and sex-matched control cell line, both at four time points (stages) (day 0, 6, 15, and 21), corresponding to the initial phase of the development of the disease. The scRNAseq data is available through the Gene Expression Omnibus (GEO), accession number GSE183248. In this study, we call a cell line at a specific time point a cell condition, leading to eight cell conditions and use a convention *cell line_day_* (e.g., Control_D0_ for control cell line at day 0; PD_D0_ for PD cell line at day 0) to refer to a particular cell condition (see Supplementary Table 1). We model the expression data of each cell condition by a matrix *E* in which rows represent genes, columns represent cells, and an entry *E_ij_*is the normalized read count of gene *i* in cell *j*.

To integrate the data with prior knowledge, we collect four molecular networks for *Homo sapiens*. To create the PPI network, we collect all physical interactions between proteins from BioGRID 4.3.195 (Oughtred *et al*., 2019), captured by at least one of the following experiments: Two-hybrid, Affinity Capture-Luminescence, Affinity Capture-MS, Affinity Capture-RNA, Affinity Capture-Western. To make the GI network, we fetch genetic interactions reported in BioGRID 4.3.195 (Oughtred *et al*., 2019). We create the COEX network by collecting the top 1% strongest correlations between genes from CoexpressDB v.7.3 (Obayashi *et al*., 2019). Finally, we construct the MI network by connecting genes participating in the same metabolic pathways in KEGG. We retrieve the pathways that are annotated by at least one of the following metabolism-related keywords in KEGG 2021/01 (Kanehisa *et al*., 2017): metabolism, metabolic, glycolysis, TCA, oxidative phosphorylation, fatty acid, pentose, degradation, or biosynthesis.

We filter the SC expression data for each cell condition to keep only protein-coding genes with at least one PPI in BioGRID, as PPIs are the most direct evidence that two proteins interact. Similarly to what was done in Malod-Dognin *et al*. (2019), we construct condition-specific PPI, GI, COEX and MI networks by considering protein-coding genes expressed in a cell condition (as measured by scRNA-seq). An edge connects nodes in the networks if the corresponding genes (or, equivalently, their protein products) interact in the molecular interaction networks obtained from the databases (detailed above) (see Supplementary Table 2).

#### 2.1.2 Biological annotations, PD genes and DEGs

To assess if our integration framework produces biologically coherent “gene embeddings”, we obtain biological annotations from Gene Ontology (GO) (Ashburner *et al*., 2000), KEGG pathways (KP) (Kanehisa *et al*., 2017) and Reactome pathways (RP) (Jassal *et al*., 2020) (all annotations were collected on 10 March 2021). We also use KPs to assess the biological relevance of our predicted set of genes and identify metabolism mechanisms perturbed during early PD development. Additionally, we collect genes from DisGeNet associated with Parkinson’s Disease (Concept Unique Identifier: C0030567) (collected on 14 May 2021) and consider them our ground-truth PD genes, terming them *DisGeNet PD genes*. We keep only those *DisGeNet PD genes* expressed in our transcriptomics data, resulting in 1,378 genes. To examine whether our methodology predicts PD-related genes that cannot be uncovered through conventional DEG analysis, we obtain the 232 protein-coding DEGs from the original study of the SC analysis Novak *et al*. (2022) and investigate their overlap with our ***Core PD predictions***.

### 2.2 NetSC-NMTF data integration model

To integrate a condition-specific single-cell expression matrix, *E*, with molecular interaction networks, we extend an NMTF-based method, iCell (Malod-Dognin *et al*., 2019), to our new framework NetSC-NMTF (see Fig. 1).

**Figure 1:**
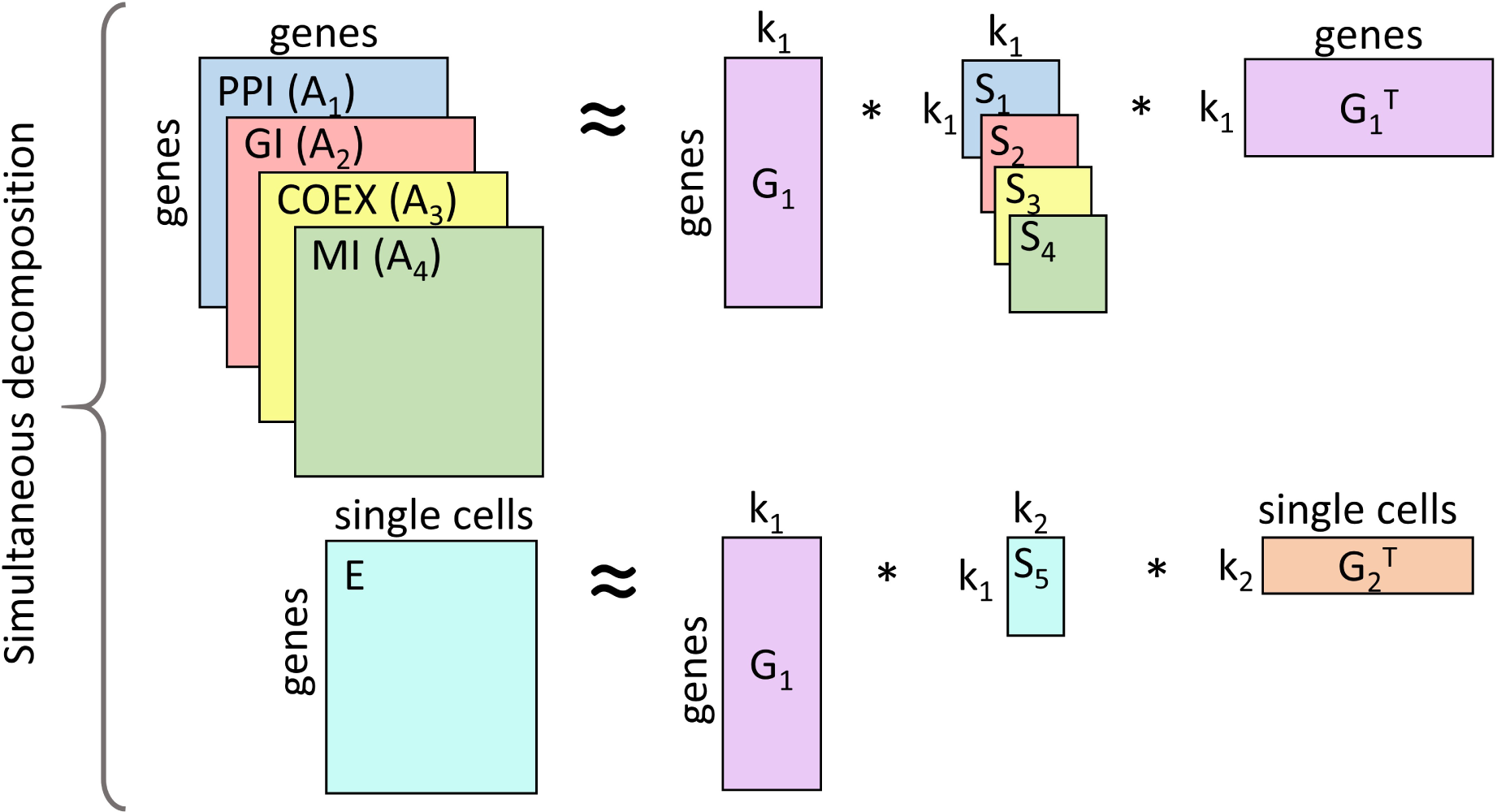
NetSC-NMTF model. Non-negative Matrix Tri-Factorization (NMTF)-based model used for integrating SC expression matrix, E, with molecular interaction networks PPI, GI, COEX, and MI (represented by their adjacency matrices, *A*_1_, *A*_2_, *A*_3_, and *A*_4_, respectively). The matrix factor *G*_1_ is shared across decompositions to allow learning from all input matrices. The parameters *k*_1_ and *k*_2_ indicate the reduced dimensions of the embedding spaces of human genes and single cells, respectively.

Molecular interaction networks are represented by their adjacency matrices, *A*_*i*∈{1,…,4}_, where each *A_i_*is a symmetric matrix with *A_i_*[*v*][*w*] entry value of one if genes *v* and *w* interact with each other, and zero otherwise. All input matrices are simultaneously decomposed into products of three matrix factors so that 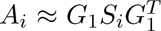 and 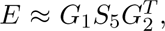 where *G*_1_ *∈* ℝ^*n*×*k*_1_^, *G*_2_ *∈* ℝ^*m*×*k_2_*^, *S*_*i*∈{1,…,4}_ ∈ ℝ^*k*_1_×*k*_1_^ and *S*_5_ *∈* ℝ^*k*_1_×*k*_2_^, with *n* and *m* being the number of genes and SCs, respectively. *k*_1_ and *k*_2_ represent the optimal dimensions of the latent embedding spaces, which we obtain by computing the maximum dispersion coefficient (Brunet *et al*., 2004) (see Supplementary Section 3.1, Supplementary Table 3 and Supplementary Figs. 2 and 3). Note that matrix factor *G*_1_ is shared across all decompositions, facilitating the information flow and learning from all data.

According to the embedding interpretation of NMTF, the set of rows of matrix *G*_1_ defines the set of embedding vectors of the genes (also called “gene embeddings”), and the set of rows of matrix *G*_2_ defines the set of embedding vectors of the SCs. To emphasize the contribution of a biological condition of SCs, we transform “gene embeddings” from *G*_1_ to the space spanned by *G*_2_ by using the transformation matrix *S*_5_ to compute *U* = *G*_1_ *∗ S*_5_ (detailed in Supplementary Section 4). In the rest of the paper, the rows of matrix *U* (that we call *u_i_*) are referred to as “gene embeddings”. On the other hand, NMTF also has a co-clustering interpretation where *G*_1_ and *G*_2_ are interpreted as cluster indicator matrices of genes and SCs, grouping genes and SCs into *k*_1_ and *k*_2_ clusters, respectively. *S*_*i*∈{1,…,4}_ matrices are interpreted as the compressed representations of molecular networks, and *S*_5_ is the compressed representation of the SC expression matrix. Based on this interpretation, we use *G*_1_ when performing clustering, as it leads to higher enrichments in biological annotations (Supplementary Fig. 4) than the clusters of matrix *U* (Supplementary Fig. 5).

Solving NMTF is an NP-hard continuous optimization problem (Vavasis, 2010). Thus, we obtain all matrix factors by applying a heuristic fixed-point solver based on multiplicative update rules (MURs) (Ding *et al*., 2006) (see Supplementary Section 1) to solve the following optimization problem:

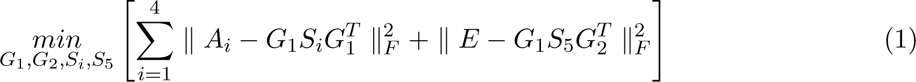

where *G*_1_*, G*_2_ *≥* 0 and ∥ ∥*_F_* denotes the Frobenius norm. Starting from an initial solution, the solver iteratively uses MURs to converge towards a locally optimal solution. To initialize the matrix factors, we apply singular value decomposition (SVD) on the original input matrices because it reduces the number of iterations needed to achieve convergence and makes the solver deterministic (Qiao, 2015). To comply with the non-negativity constraint of NMTF, we generate the initial solution by taking the absolute values of the entries of the resulting SVD matrices. The iterative process stops when the objective function converges (e.g., Supplementary Fig. 1), which we measure every 10 iterations with 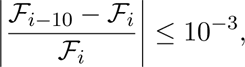 where *ℱ_i_* is the value of the objective function at the current iteration and *ℱ^i^*^−10^ is the value computed at iteration *i* − 10.

### 2.3 Definition of the “gene movement”

To determine how different biological states/conditions of SCs alter “gene embeddings”, we introduce the so-called *Gene Mapping Matrix* (*GMM*). *GMM* is a symmetric distance matrix that captures the relative positions between the “gene embeddings” in the embedding space of SCs of one cell condition. We compute a *GMM* of a cell condition with *GMM* [*u_i_*][*u_j_*] = *d*(*u_i_, u_j_*)/∥*U* ∥*_F_*, where each entry corresponds to the norm-scaled Euclidean distance between the “gene embeddings” *u_i_* and *u_j_* of two genes *i* and *j* in the matrix *U* and ∥ ∥*_F_* denotes the Frobenius norm. Since *GMMs* encode such relative and normalized gene positions, we can directly compare the position of one gene in one *GMM*, *g_i_*, with its position in another *GMM*, 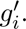 To do this, we compute the Euclidean distance between these positions with 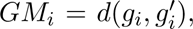 where *GM_i_* is a scalar that we call a “movement” of a gene *i* between two cell conditions (i.e., “gene movement”). In other words, “gene movement” represents how the relative position of a gene (i.e., relative to all other genes in one cell condition) changes between the embedding spaces of two cell conditions. While “gene movement” can be computed for genes between any two cell conditions, we calculate the “gene movement” of genes between time point-matching PD and control cell conditions in our study. This allows us to investigate how PD affects the spatial organization of embedding spaces of genes compared to healthy controls at different stages of PD development. The “gene movement” is either a positive value and indicates to what extent PD alters the relative position of a gene compared to a corresponding control, or zero if there is no such change.

### 2.4 Predicting novel PD-associated genes: A 2-step downstream method

We uncover novel PD-associated genes by applying our NetSC-NMTF framework to obtain “gene embeddings” of time point-specific data, which we mine using the following 2-step downstream method.

In the first step, we obtain PD predictions for a given time point (i.e., ***Stage-specific PD predictions***). This step is based on the hypothesis that *DisGeNet PD genes* (defined in Section 2.1.2) cluster in the gene embedding spaces of PD cell conditions (see Section 3.1) and that genes that group with *DisGeNet PD genes* are also relevant for PD. A similar approach has been successfully applied by Gligorijević *et al*. (2016) to find new cancer driver genes. Therefore, we apply a k-means clustering algorithm (Pedregosa *et al*. (2011); a widely used algorithm for getting gene clusters) on *G*_1_ matrix of a time point-specific PD cell condition. Next, we perform an enrichment analysis (see Supplementary Section 2) in *DisGeNet PD genes* of these clusters and keep the significantly enriched ones. Finally, from significantly enriched clusters, we retain those genes that are not labeled as *DisGeNet PD genes* and are also expressed in the control cells of the corresponding time point, calling this set of genes ***Stage-specific PD predictions***. We apply the first step for each of the four time points, resulting in four sets of ***Stage-specific PD predictions***, which we validate in the literature in Supplementary Section 5, Supplementary Figs. 9 and 10.

In the second step, we intersect the four ***Stage-specific PD predictions*** obtained from step 1 to define our ***Core PD predictions***. By focusing on the intersection of ***Stage-specific PD predictions***, we remove the potential stage-specific noise and hypothesize that we uncover genes that drive PD across all time points caused by the *PINK1* mutation. Furthermore, previous studies on time-dependent SC data have shown that a significant signal could be detected when SC data across all time points are exploited, stressing the importance of such approaches (Jung *et al*., 2020; Novak *et al*., 2022). To prioritize the ***Core PD predictions***, we compute their average “movement” across all time points (Section 2.3) and rank the predictions according to their average “movement”, the largest first. This approach is based on the observation that the relative positions of *DisGeNet PD genes* are more altered in PD versus control conditions compared to other genes (see Section 3.1). Therefore, we hypothesize that the more the relative position of a gene is altered in PD versus control (at a given time point, or across all time points), the more relevant it is for PD (which we confirm in Supplementary Section 5 and Supplementary Fig. 8).

### 2.5 Validating predictions

We validate our gene predictions and quantify their association with PD in the literature by using an automated PubMed publication search to count the co-occurrence of each gene from the set of predictions and the background set (defined below) with the term “Parkinson’s disease” in PubMed publications. To measure if our predicted genes are significantly more co-occurring with PD in the literature, we perform a one-sided Mann-Whitney U (MWU) test (with a significance level of 5%) between the co-occurrence distributions of the set of predictions and the corresponding background. For validating each set of ***Stage-specific PD predictions***, we define the background set of genes as genes that are expressed at a particular time point and remove *DisGeNet PD genes* and the set of ***Stage-specific PD predictions***. For validating the ***Core PD predictions***, we define the background set of genes as genes that are expressed across all time points and do not belong to the *DisGeNet PD genes* and the ***Core PD predictions***.

For a complementary validation of our predictions, we do an additional validation experiment evaluating if our sets of gene predictions are statistically significantly related to PD. We perform enrichment analysis (Supplementary Section 2) (significance threshold of 0.05) in a set of PD-related genes less validated in the literature than *DisGeNet PD genes* on each set of our gene predictions against the background (defined above). The set of PD-related genes are genes that: 1) co-occur with the term “Parkinson’s disease” in at least one PubMed study, in an automatic literature search of PubMed (2,031 genes) (search was performed on 13 May 2022), or 2) are in the Gene4PD database (a database containing PD associated genes extracted from more than 60 genomic data sources) (2,121 genes) (Li *et al*., 2021) (collected on 07 October 2021). The overlap between the two sets is 517 genes, resulting in 3,635 unique PD-related genes.

## 3 Results and Discussion

We apply our NetSC-NMTF data integration framework (see Section 2.2) to scRNA-seq timeseries data and molecular networks to obtain “gene embeddings”, which we investigate with our 2-step downstream mining method (see Section 2.4) to uncover impaired PD pathways, novel PD-associated genes and suggest drug-repurposing options.

### 3.1 *DisGeNet PD genes* have specific properties in the embedding spaces of genes and single cells

Integrating SC expression data with prior knowledge in all four molecular networks produces functionally meaningful “gene embeddings”, as measured by the clustering and enrichment analysis in GO, KP and RP annotations (Supplementary Section 3.2). We obtain gene clusters by applying the k-means clustering algorithm (Pedregosa *et al*., 2011) (a widely used algorithm for getting gene clusters) on the “gene embeddings” of each time point-specific cell condition. Consequently, we see that genes embedded close to each other are functionally related, leading to the interpretation that such genes participate in the same molecular pathways. As PD is characterized by perturbation of many molecular mechanisms, we assume that *DisGeNet PD genes* participate in the same molecular pathways and investigate if *DisGeNet PD genes* are also embedded close to each other. Therefore, we perform an enrichment analysis (detailed in Supplementary Section 2) in *DisGeNet PD genes* of the clusters described above, measuring the percentage of clusters significantly enriched in *DisGeNet PD genes* (*p-value ≤* 5%). We observe around 18% of significantly enriched clusters across all cell conditions (see Supplementary Fig. 6), where approximately half of the genes in these clusters are *DisGeNet PD genes* (average fold enrichment is 2.06). These results indicate that *DisGeNet PD genes* are not interspersed throughout the gene embedding spaces but rather group together.

Furthermore, we hypothesize that *DisGeNet PD genes* participate in molecular pathways that are altered more than the pathways characterized by other expressed genes between control and PD cell conditions at individual time points. To measure this alteration, we apply the method described in Section 2.3 to compute the distributions of the changes in the relative position (i.e., “gene movement”) of *DisGeNet PD genes* and non-*DisGeNet PD genes* (background) between the embedding spaces of each PD cell condition and its time point-matching control. We compare the two “gene movement’ distributions at each time point by performing a one-sided Mann-Whitney U (MWU) test (with a significance level of 5%). The *DisGeNet PD genes* “movement” distributions across all time points are statistically significantly larger than the one of background genes, with *p-values ≤* 1.65*e^−^*^05^ (see Supplementary Fig. 7).

In conclusion, we observe that: 1) *DisGeNet PD genes* cluster together in the gene embedding spaces of individual cell conditions; 2) the “movement” of *DisGeNet PD genes* is statistically significantly larger than that of background genes for each time point-specific control and PD pairwise comparison. The following sections build upon these two observations to predict and validate novel PD-associated genes.

### 3.2 Obtaining Core PD predictions

To uncover novel PD-associated genes, we mine the “gene embeddings” obtained with our NetSCNMTF framework with our 2-step downstream method (see Section 2.4). In the first step, obtain PD predictions for a given time point (i.e., ***Stage-specific PD predictions***), by extracting non-*DisGeNet PD genes* from the clusters of PD cell conditions (obtained above) significantly enriched in *DisGeNet PD genes*. We show that relevant biological information can be gained from analyzing SC data at individual time points by observing that gene predictions associated with all four time points of cell development (i.e., ***Stage-specific PD predictions***) are significantly associated with PD (see Supplementary Section 5, Supplementary Figs. 9 and 10).

To account for the noisiness of the scRNA-seq measurements and to focus on the genes involved in the PD progression across all time points, we adopt a consensus approach used in other studies (Novak *et al*., 2022; Jung *et al*., 2020) that makes a final list of predictions based on all available time points. Thus, we apply the second step of our 2-step downstream method to define a final set of predictions by intersecting all sets of ***Stage-specific PD predictions*** and ranking the genes in the overlap according to the average “movement” across all time points so that the genes with the largest average “movement” are ranked the highest (Section 2.4). This results in 193 PD predictions that we name ***Core PD predictions*** (Supplementary File 1). To assess if such a consensus approach is applicable to extract novel PD-associated genes, we test if the four sets of ***Stage-specific PD predictions*** significantly overlap by applying a sampling with replacement technique (Supplementary Section 6). Over 10,000 repetitions, the overlap of ***Stage-specific PD prediction*** sets is always larger compared to random (*p-value ≤* 1*e^−^*^05^). This significant overlap shows that a consensus approach is possible and can be used to uncover a set of new genes predicted to be related to PD across all time points.

### 3.3 *Core PD predictions* are relevant for PD

To determine the PD-relevance of the ***Core PD predictions***, we follow the validation procedures described in Section 2.5. We obtain the co-occurrence distributions of the predictions and the background genes with the term “Parkinson’s disease” in PubMed publications, which we compare with a one-sided MWU test (with a significance level of 5%). We find that the co-occurrence distribution of ***Core PD predictions*** is significantly greater than the one of background (*p-value* = 4.96*e^−^*^12^) (see Supplementary Fig. 11). Furthermore, we observe that our ***Core PD predictions*** are significantly enriched in the PD-related genes (*p-value* = 1.48*e^−^*^10^), with 49.7% of our ***Core PD predictions*** belonging to the set of PD-related genes. These results demonstrate that the 193 ***Core PD predictions*** are significantly associated with PD.

We also examine whether our methodology predicts PD-related genes that can not be uncovered through conventional DEG analysis. Thus, we compare our ***Core PD predictions*** with the 232 protein-coding DEG-based predictions obtained by Novak *et al*. (2022) (Section 2.1.2) (note that the same scRNA-seq dataset is used in our study), by checking the overlap between the two sets of genes. We uncover eight genes (*GOLT1B*, *PDIA6*, *RPN2*, *PFKP*, *FOS*, *EGLN3*, *GNAS* and *LMAN1*) (Fig. 2, a), all of which have been linked with PD (Supplementary Table 4), but whose exact roles in the pathogenesis of PD are not fully understood. Identifying these genes through two independent analyses suggests their importance in PD and warrants more extensive studies to elucidate their involvement in PD and investigate them as potential intervention opportunities. Additionally, by integrating scRNA-seq data with prior knowledge, we reduce noise characteristic for the scRNA-seq measurements, which allows us to predict and prioritize a set of PD-related genes beyond standard DEG analysis, offering new insights into PD and further proving the value of our data integration model and the 2-step downstream analysis.

**Figure 2:**
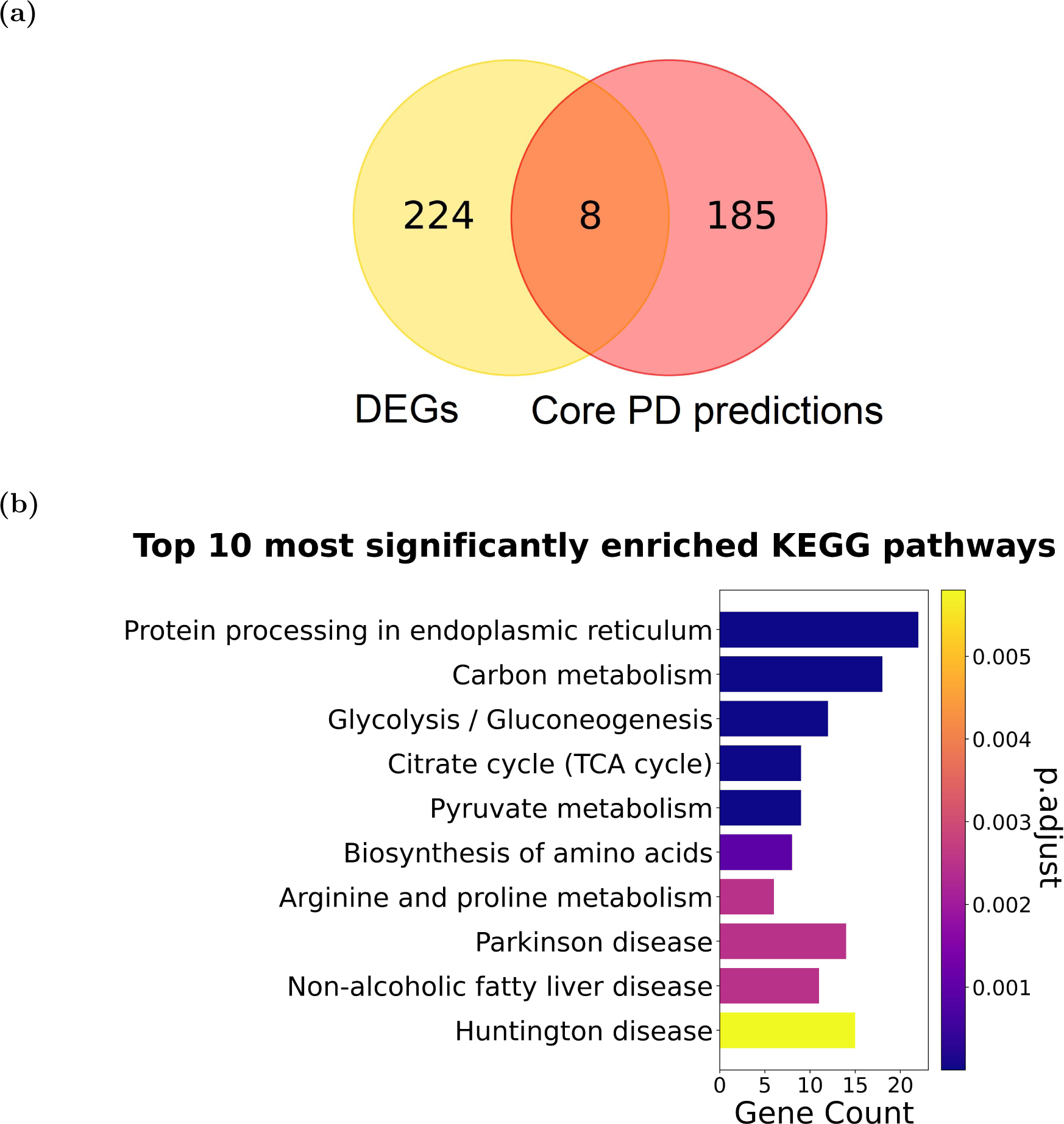
**(a) Overlap between DEGs from Novak *et al*. (2022) and *Core PD predictions*. (b) Top 10 KPs significantly enriched in *Core PD predictions***. p.adjust represents adjusted *p-values* (obtained from enrichment analysis) for multiple hypothesis testing using Benjamini and Hochberg (1995). Gene Count are the number of *Core PD predictions* that participate in a KP.

In conclusion, we prove the PD-relevance of ***Core PD predictions*** and show that the consensus approach allows for uncovering novel PD-associated genes. Furthermore, in Supplementary Section 7, we show that our 2-step method outperforms other approaches based on: 1) DEGs (LIGER (Welch *et al*., 2019), a state-of-the-art method for integrating SC data), 2) distance of non-*DisGeNet PD genes* to *DisGeNet PD genes* in the embedding spaces of PD cell conditions, or 3) “movement” of genes between control and PD time point-specific cell conditions. Our method leads to more PD-relevant predictions, which we measure by validating the predictions obtained with other methods through an automated PubMed search, and performing an enrichment analysis in a PD-related set of genes (Section 2.1.2) and KPs.

### 3.4 KEGG pathways enriched in our *Core PD predictions* are associated with PD

The disruption of many molecular pathways characterizes PD, indicating that PD is a metabolic disorder (Anandhan *et al*., 2017). Thus, we determine the metabolic functions associated with our ***Core PD predictions*** and investigate if they agree with the literature. To that aim, we perform enrichment analysis (Supplementary Section 2) in KPs, identifying 37 significantly enriched ones (Supplementary File 2).

Here, we present the top 10 most significantly enriched KPs of the ***Core PD predictions*** (see Fig. 2, b), which we rank according to the most significant *p-value* (smallest first). Most importantly, *Parkinson’s disease* is one of the most enriched pathways (rank 8), further showing that our predictions are related to this disease. *Protein processing in the endoplasmic reticulum* (ER) is the first-ranked pathway. It is indeed relevant for PD as the accumulation of *α*-synuclein proteins in PD induces ER stress by inhibiting the ER-Golgi trafficking and leading to the dysregulation of protein processing in ER and eventually cell death (Ghemrawi and Khair, 2020). *Carbon metabolism* (rank 2) is altered in PD, leading to a decrease in glucose metabolism and abnormally elevated levels of pyruvate (*pyruvate metabolism*, rank 5) (Anandhan *et al*., 2017). Furthermore, *glycolysis/gluconeogenesis* (rank 3) and *citrate cycle* (rank 4) contribute to the overarching *carbon metabolism pathway* (Kanehisa *et al*., 2017), emphasizing the contribution of the altered carbon metabolism in the progression of PD. Thus, protein processing in the ER and carbon metabolism undoubtedly play a vital role in the progression of PD and should be further investigated. PD is also characterized by the dysregulation of the *biosynthesis of amino acids* (rank 6) (Dong *et al*., 2022), with evidence pointing to a lower abundance of arginine and unaltered amounts of proline (*arginine and proline metabolism*, rank 7) (Figura *et al*., 2018), highlighting the need to study these mechanisms in more detail and uncover their exact role in PD. Although the association between the *non-alcoholic fatty liver disease pathway* (rank 9) and PD is not well understood, evidence suggests that it plays a role in PD (Chi *et al*., 2018). Finally, it is not surprising that Huntington’s disease (rank 10) is one of the most enriched pathways as it shares many disrupted mechanisms with PD, including protein processing in ER (Roussel *et al*., 2013).

Overall, we show that the top 10 KPs associated with our ***Core PD predictions*** are all relevant for PD development at the early stages of the disease. Additionally, we perform enrichment analysis in KPs of the 232 DEGs from Novak *et al*. (2022), revealing that they are not enriched in any pathway, further showing that our methodology uncovers PD-related genes that complement the findings obtained by the traditional DEG approach. In Supplementary Section 8, we also perform an enrichment analysis in KPs of *DisGeNet PD genes* and compare the results with those of ***Core PD predictions***. Notably, we find that *DisGeNet PD genes* are enriched in 144 KPs, sharing 14 common pathways with the ***Core PD predictions*** that most importantly include *Parkinson’s disease* (rank 8). Thus, we suggest that future research endeavors aim to further understand the relationship of the discussed pathways with PD, as they might significantly contribute to the pathogenesis of this disease and provide opportunities for drug discovery efforts.

### 3.5 Literature validation of the top 20 *Core PD predictions*

The results above show that our ***Core PD predictions*** are globally associated with PD, suggesting that the remaining non-validated ***Core PD predictions*** are also relevant for PD. Thus, we focus on the top 20 prioritized ***Core PD predictions*** to better characterize their relationship to PD and their potential role in drug-repurposing strategies. We manually assess if the top 20 ***Core PD predictions*** are related to PD in the literature and find that eight genes (40%) have a known association. We find literature evidence for the remaining 12 genes that could explain their potential role in the disease (Supplementary Table 5). Additionally, we identify seven (out of 12) druggable genes that represent candidates for future drug-repurposing investigations and suggest two other genes for drug discovery studies, providing potential novel therapeutic opportunities for PD (Supplementary Table 4). The 12 gene predictions could play a role in PD based on the metabolic pathways they participate in, or their involvement in other neurodegenerative diseases. Here, we discuss seven of those predictions and their druggability.

The mutation of *PFN1* (rank one) leads to the development of a neurodegenerative disease called amyotrophic lateral sclerosis (Teyssou *et al*., 2022). *PFN1* also regulates the dynamics of the actin cytoskeleton, whose dysregulation has been implicated in multiple neurodegenerative diseases such as PD and Alzheimer’s. *PFN1* is also a target of Artenimol, a drug originally used to treat malaria (Organization, 2015). Gao *et al*. (2020) proved that Artenimol could be used for treating neuroinflammatory diseases by inactivating the PI3K/AKT and NF-*κ*B signalling pathways, two pathways that are dysregulated in PD (Rai *et al*., 2019; Singh *et al*., 2020), suggesting that the Artenimol-*PFN1* drug-target interaction could be exploited for treating PD. We highlight six gene predictions (*APLP2*, *RRBP1*, *RCN1*, *SEC63*, *KDELR1*, *SSR4*) for their role in maintaining the proper functioning of ER (see Supplementary Table 4). *APLP2* (rank 4) is a target of zinc and some of its compounds. It could be exploited for maintaining the optimal levels of zinc, whose alterations have been implicated in the pathophysiology of PD (Sikora and Ouagazzal, 2021). We also find that *RRBP1* (rank 5) is druggable by Radezolid (Wishart *et al*., 2018), which has been used in trials to treat skin diseases and might be repurposed for PD. *RCN1* (rank 8) and *SSR4* (rank 15) are affected by calcium (Wishart *et al*., 2022), providing opportunities to maintain *Ca*^2+^ homeostasis and reverse the toxicity of the misfolded *α*-synuclein proteins, thereby preventing ER stress. Given that four of the six genes under investigation are established drug targets, the remaining two, *SEC63* and *KDELR1*, warrant consideration as prospective candidates for future PD drug discovery studies.

Here, we have shown evidence that the top prioritized ***Core PD predictions*** are associated with PD, motivating us to propose 12 novel PD-associated genes. Seven of the 12 uncovered genes are known drug-targets and candidates for future drug-repurposing investigations, while two genes represent possible intervention points for drug discovery studies.

### 3.6 *Core PD predictions* are associated with PD subtype carrying a *PINK1* mutation

To determine if ***Core PD predictions*** are relevant to the *PINK1* subtype of PD considered in this study, we investigate if our gene predictions are closely related to the *PINK1* gene by studying the subgraph that the predictions and *PINK1* form in the PPI network. The intuition behind this approach is that proteins in the same neighbourhood in a PPI network are likely to participate in the same functional modules, such as protein complexes, metabolic pathways or signal transduction systems. We generate the subgraph of the PPI network obtained from Biogrid (defined in Section 2.1) induced by the genes expressed in at least one of our cell conditions, making it more relevant for our data. In this data-specific PPI network, we measure the shortest path of ***Core PD predictions*** and background (genes in the PPI subgraph that are not ***Core PD predictions***) to the *PINK1* gene and compare the two shortest paths distributions using a one-sided MWU test (with a significance level of 5%). We observe that ***Core PD predictions*** are statically significantly closer (*p-value* = 6.91*e^−^*^17^) to the *PINK1* gene than the background (see Fig. 3, a), with the average shortest path length of 1.92 of our predictions and 2.26 of the background. To determine if our predictions are more specific to the *PINK1* subtype of PD than the 232 DEGs from Novak *et al*. (2022), we perform the same experiment with this set of genes and observe that the DEGs are also statistically significantly closer (*p-value* = 2.06*e^−^*^04^) to the *PINK1* gene than the background (genes in the PPI subgraph that are not DEGs). However, as the average shortest path of DEGs to *PINK1* is larger than that of our ***Core PD predictions*** (2.13 compared to 1.92), we conclude that our method allows us to find genes that are more specific to the *PINK1* subtype of PD.

**Figure 3:**
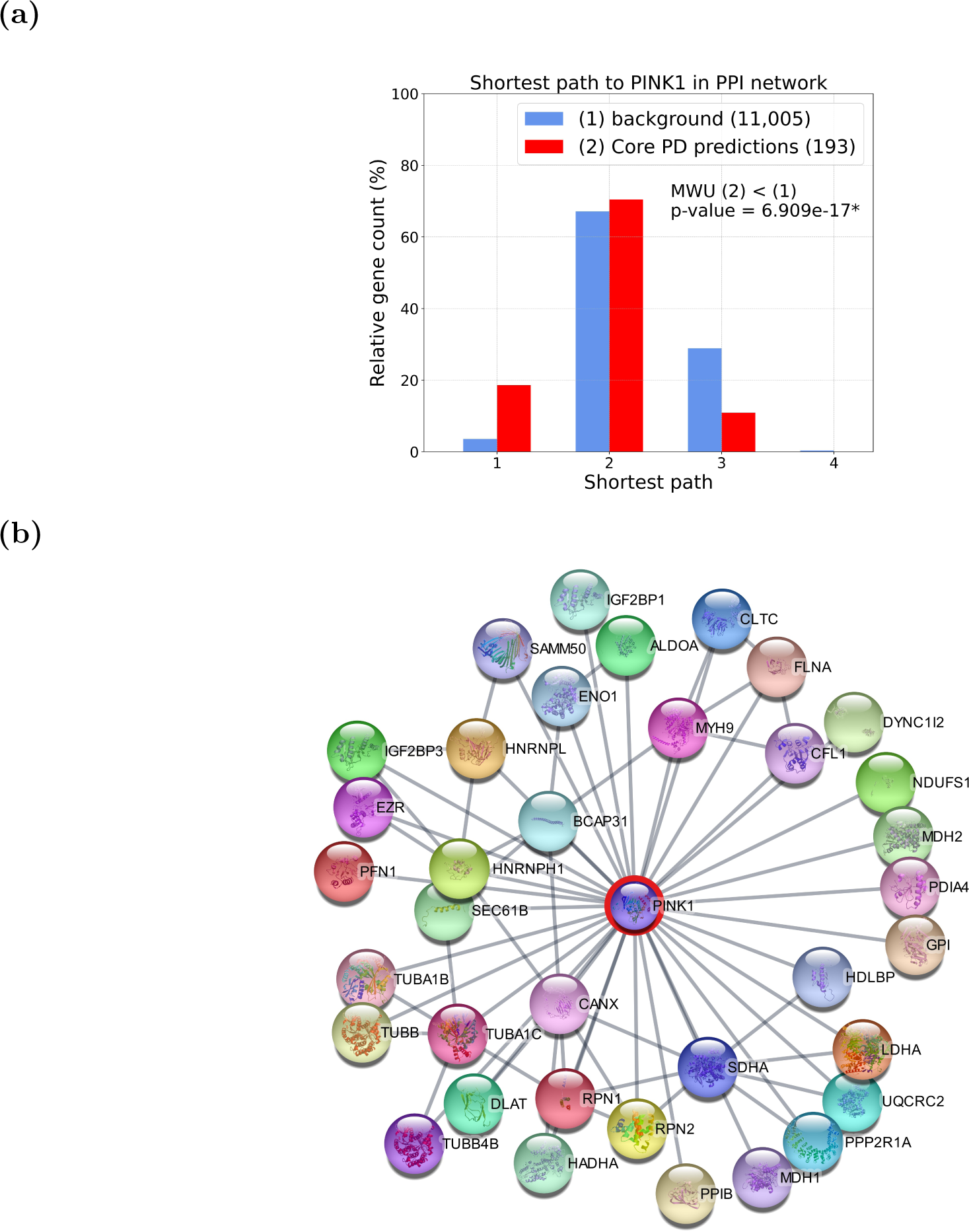
**(a) Shortest path length distribution of genes to *PINK1* gene in the PPI network**. We represent the background gene set as group (1) (blue) and ***Core PD predictions*** set as group (2) (red), indicating the number of genes in each set in the brackets on the right. MWU (2) *<* (1) indicates that we perform a one-sided MWU (with a significance level of 0.05) to test if the shortest path distribution of ***Core PD predictions*** is significantly smaller than the one of background genes (with *p-value <* 0.05 indicated by *). **(b) PPI network of *PINK1* and its 36 first neighbours from *Core PD predictions*.** Network visualization was done using Cytoscape 3.10.0 (Shannon *et al*., 2003).

Having shown that ***Core PD predictions*** are close to *PINK1* in the PPI network, we hypothesize that they participate in the same PD-related pathways with *PINK1*. We further assume that the genes closest to *PINK1* in the PPI network are the ones that experience the effects of *PINK1* ’s mutation first, subsequently leading to the impairment of the biological mechanisms these genes participate in, thereby contributing to PD development. Therefore, we focus on the 36 ***Core PD predictions*** that are *PINK1* ’s first network neighbours (see Fig. 3, b). To test if these genes indeed participate in PD-associated biological mechanisms and how they relate with *PINK1*, we perform an enrichment analysis in PD terms (i.e., pathways) obtained from PD map (Fujita *et al*., 2014); we exclude *Scrapbook* and *Parkinsons UK Gene Ontology genes* from the terms collected from PD map, as they do not represent biological pathways. We find that our 36 ***Core PD predictions*** are statistically significantly enriched in *glycolysis* (*p-value* = 5.68*e^−^*^03^), *actin filament organization* (*p-value* = 9.07*e^−^*^03^), *mitochondrial and ROS metabolism* pathway (*p-value* = 4.71*e^−^*^02^), and *axonal remodeling and CDK5 signaling* (*p-value* = 3.10*e^−^*^02^). A recent study (Travaglio *et al*., 2023) showed that glycolysis is elevated in a PD model harbouring a *PINK1* I368N mutation, a mutation also investigated in this paper. Additionally, altered actin dynamics were observed in *PINK1* knockdown dopamine neuronal cells (Kim and Son, 2010). Mitochondrial dysfunction and increased mitochondrial ROS are known as hallmarks of the PD subtype carrying *PINK1* mutations (Requejo-Aguilar *et al*., 2014). Interestingly, there is no direct evidence linking *PINK1* with *axonal remodelling and CDK5 signaling* mechanism. However, *PINK1* mutations influence *LLRK2* levels (Azkona *et al*., 2018) (a commonly mutated protein in PD) (Shah and Rossie, 2018), which in turn disrupts *axonal remodelling and CDK5 signaling* in PD. Therefore, *PINK1* mutations may contribute to PD development by disrupting *axonal remodelling and CDK5 signaling* through their relationships with *LLRK2*. Our findings emphasize the need to investigate *axonal remodelling and CDK5 signaling* as a new and important pathway in PD pathogenesis associated with *PINK1* mutations.

These results demonstrate the power of our methodological pipeline to predict genes pertinent to a PD subtype characterized by a *PINK1* mutation. Additionally, the PPI subnetwork from Fig. 3, b) reveals the molecular interactions that connect the *PINK1* mutations with the four PD pathways mentioned above. To further understand how a mutation in *PINK1* leads to PD development, we recommend future research to be directed at studying the 36 ***Core PD predictions*** that are *PINK1* ’s first PPI network neighbours and their related molecular mechanisms, such as *axonal remodelling and CDK5 signaling*.

## 4 Conclusion

The complexity of PD requires new integration methods capable of exploiting multi-omics data. To address the open challenge of integrating time-series scRNA-seq data with molecular networks to uncover novel PD-associated genes beyond DEGs, we propose our NetSC-NMTF framework (Section 2.2) and the 2-step downstream method (Section 2.4). Using prior knowledge hidden in the molecular networks, our framework effectively minimizes noise intrinsic to the scRNA-seq measurements, thereby prioritizing genes and pathways pertinent to PD.

We apply our methodology to integrate and analyze four condition-specific molecular interaction networks and scRNA-seq data of a cell line harbouring a PD-associated mutation in the *PINK1* gene (I368N), or a control cell line, at four time points of cell development (eight cell conditions). We identify 193 PD-related predictions that we call ***Core PD predictions***, out of which 49.7% have been previously associated with PD in the literature. We also show that DEG-based approaches cannot uncover our genes and that our method predicts genes that are more PD relevant than DEG-based methods such as LIGER (Welch *et al*., 2019) and the method presented in Novak *et al*. (2022) (Section 3.3, Fig. 2, a)). To shed light on biologically relevant PD mechanisms, we discuss the top 10 most enriched KPs of the ***Core PD predictions*** (see Fig. 2, b). We perform a manual literature validation on the top 20 predictions to suggest 12 novel PD-associated genes implicated in PD based on the metabolic pathways they participate in, or their role in other neurodegenerative disorders. Seven out of 12 novel PD-associated genes are druggable, so we recommend future drugrepurposing directions that could represent new therapeutic options for treating PD. Additionally, we predict two new PD-associated genes that are not known drug targets but represent potential intervention points that should be considered in future drug discovery studies on PD. Furthermore, we demonstrate that the ***Core PD predictions*** are specific for the PD subtype associated with the *PINK1* gene mutation. To our knowledge, this is the only non-DEG-based approach that integrates scRNA-seq time-series data of Parkinson’s disease and control samples with molecular networks and exploits the information of the “gene embeddings” to: 1) uncover novel disease-related genes, 2) reveal critical metabolic pathways and 3) propose new drug-repurposing and drug discovery options.

Here, we analyze iPSC-derived data from a mutation-specific neuronal cell line against the control. While the iPSC-derived neuronal models represent the gold standard for analyzing PD in vitro, significantly contributing to understanding this disease (Novak *et al*., 2022; Smajić *et al*., 2022), it suffers from several biological drawbacks. For example, ageing-related effects and epigenetic influence are lost during reprogramming to iPSCs. Additionally, as iPSC is a two-dimensional cell culture model, it does not fully recapitulate the cell-cell/cell-matrix interactions and cell morphology present in vivo (Lopes *et al*., 2017). However, our methodology is generic and could be modified to accommodate data from tissue samples and other data types, such as metabolomics, proteomics, and epigenetics (including bulk datasets). New data could complement the information in the expression data and molecular networks, allowing for uncovering novel biological knowledge. Finally, our framework could also be applied to other PD mutations, diseases, or processes where analysis of time series expression data is key, e.g., gender ageing differences, or cell response to drug treatment.

## Author Contributions

Conceptualization, N.P. and N.M.D.; Methodology, N.P., N.M.D., G.C. and K.M.; Validation, K.M.; Formal Analysis, K.M.; Investigation, K.M.; Resources, N.P., A.S., G.N. and D.K.; Writing – Original Draft, G.C. and K.M.; Writing – Review & Editing, N.P., A.S., N.M.D. and K.M.; Visualization, K.M.; Supervision, N.P.; Funding Acquisition, N.P. and A.S.

## Code Availability

The source code of NetSC-NMTF, the subsequent 2-step mining method, a step-by-step tutorial for applying the methodological framework and the data used in this study can be accessed at https://github.com/KatarinaMihajlovic/NetSCNMTF-2stepmining.

## References

Anandhan, A. et al. (2017). Metabolic dysfunction in parkinson’s disease: bioenergetics, redox homeostasis and central carbon metabolism. Brain Research Bulletin, 133, 12–30.

Argelaguet, R. et al. (2020). Mofa+: a statistical framework for comprehensive integration of multi-modal single-cell data. Genome Biology, 21(1), 1–17.

Ashburner, M. et al. (2000). Gene ontology: tool for the unification of biology. Nature Genetics, 25(1), 25–29.

Azkona, G. et al. (2018). Lrrk2 expression is deregulated in fibroblasts and neurons from parkinson patients with mutations in pink1. Molecular Neurobiology, 55(1), 506–516.

Balestrino, R. and Schapira, A. (2020). Parkinson disease. European Journal of Neurology, 27(1), 27–42.

Benjamini, Y. and Hochberg, Y. (1995). Controlling the false discovery rate: a practical and powerful approach to multiple testing. Journal of the Royal Statistical Society: series B (Methodological), 57(1), 289–300.

Brunet, J.-P. et al. (2004). Metagenes and molecular pattern discovery using matrix factorization. Proceedings of the National Academy of Sciences, 101(12), 4164–4169.

Chi, J. et al. (2018). Integrated analysis and identification of novel biomarkers in parkinson’s disease. Frontiers in Aging Neuroscience, 10, 178.

Ding, C. et al. (2006). Orthogonal nonnegative matrix t-factorizations for clustering. In Proceedings of the 12th ACM SIGKDD International Conference on Knowledge Discovery and Data mining, pages 126–135.

Dong, C. et al. (2022). Plasma metabolite signature classifies male lrrk2 parkinson’s disease patients. Metabolites, 12(2), 149.

Elyanow, R. et al. (2020). netnmf-sc: leveraging gene–gene interactions for imputation and dimensionality reduction in single-cell expression analysis. Genome Research, 30(2), 195–204.

Figura, M. et al. (2018). Serum amino acid profile in patients with parkinson’s disease. PLoS One, 13(1), e0191670.

Fujita, K. A. et al. (2014). Integrating pathways of parkinson’s disease in a molecular interaction map. Molecular Neurobiology, 49, 88–102.

Gao, Y. et al. (2020). Dihydroartemisinin ameliorates lps-induced neuroinflammation by inhibiting the pi3k/akt pathway. Metabolic Brain Disease, 35(4), 661–672.

Ghemrawi, R. and Khair, M. (2020). Endoplasmic reticulum stress and unfolded protein response in neurodegenerative diseases. International journal of molecular sciences, 21(17), 6127.

Giguère, N., et al. (2018). On cell loss and selective vulnerability of neuronal populations in parkinson’s disease. Frontiers in Neurology, page 455.

Gligorijević, V., et al. (2016). Patient-specific data fusion for cancer stratification and personalised treatment. In Biocomputing 2016: Proceedings of the Pacific Symposium, pages 321–332. World Scientific.

Huizing, G.-J. et al. (2023). Paired single-cell multi-omics data integration with mowgli. bioRxiv, pages 2023–02.

Jassal, B. et al. (2020). The reactome pathway knowledgebase. Nucleic Acids Research, 48(D1), D498–D503.

Jung, I. et al. (2020). A non-negative matrix factorization-based framework for the analysis of multi-class time-series single-cell rna-seq data. IEEE Access, 8, 42342–42348.

Kanehisa, M. et al. (2017). Kegg: new perspectives on genomes, pathways, diseases and drugs. Nucleic Acids Research, 45(D1), D353–D361.

Kim, K.-H. and Son, J. H. (2010). Pink1 gene knockdown leads to increased binding of parkin with actin filament. Neuroscience Letters, 468(3), 272–276.

Kunes, R. Z. et al. (2023). Supervised discovery of interpretable gene programs from single-cell data. Nature Biotechnology, pages 1–12.

Lee, J. et al. (2020). Single-cell multiomics: technologies and data analysis methods. Experimental & Molecular Medicine, 52(9), 1428–1442.

Li, B. et al. (2021). Gene4pd: A comprehensive genetic database of parkinson’s disease. Frontiers in Neuroscience, 15.

Li, M. M. et al. (2023). Contextualizing protein representations using deep learning on protein networks and single-cell data. bioRxiv.

Lopes, F. M. et al. (2017). Mimicking parkinson’s disease in a dish: merits and pitfalls of the most commonly used dopaminergic in vitro models. NeuroMolecular Medicine, 19, 241–255.

Luecken, M. D. and Theis, F. J. (2019). Current best practices in single-cell rna-seq analysis: a tutorial. Molecular Systems Biology, 15(6), e8746.

Luecken, M. D. et al. (2022). Benchmarking atlas-level data integration in single-cell genomics. Nature Methods, 19(1), 41–50.

Malod-Dognin, N. et al. (2019). Towards a data-integrated cell. Nature Communications, 10(1), 1–13.

Novak, G. et al. (2022). Single-cell transcriptomics of human ipsc differentiation dynamics reveal a core molecular network of parkinson’s disease. Communications Biology, 5(1), 1–19.

Obayashi, T. et al. (2019). Coxpresdb v7: a gene coexpression database for 11 animal species supported by 23 coexpression platforms for technical evaluation and evolutionary inference. Nucleic Acids Research, 47(D1), D55– D62.

Organization, W. H. (2015). Guidelines for the treatment of malaria. World Health Organization.

Oughtred, R. et al. (2019). The biogrid interaction database: 2019 update. Nucleic Acids Research, 47(D1), D529– D541.

Pedregosa, F. et al. (2011). Scikit-learn: Machine learning in python. the Journal of machine Learning research, 12, 2825–2830.

Peng, W. et al. (2019). Predicting protein functions through non-negative matrix factorization regularized by proteinprotein interaction network and gene functional information. In 2019 IEEE International Conference on Bioinformatics and Biomedicine, pages 86–89. IEEE.

Poewe, W. et al. (2017). Parkinson disease. Nature Reviews Disease Primers, 3(1), 1–21.

Qiao, H. (2015). New svd based initialization strategy for non-negative matrix factorization. Pattern Recognition Letters, 63, 71–77.

Rai, S. N. et al. (2019). The role of pi3k/akt and erk in neurodegenerative disorders. Neurotoxicity Research, 35(3), 775–795.

Requejo-Aguilar, R. et al. (2014). Pink1 deficiency sustains cell proliferation by reprogramming glucose metabolism through hif1. Nature Communications, 5(1), 4514.

Roussel, B. D. et al. (2013). Endoplasmic reticulum dysfunction in neurological disease. The Lancet Neurology, 12(1), 105–118.

Shah, K. and Rossie, S. (2018). Tale of the good and the bad cdk5: Remodeling of the actin cytoskeleton in the brain. Molecular Neurobiology, 55, 3426–3438.

Shannon, P. et al. (2003). Cytoscape: a software environment for integrated models of biomolecular interaction networks. Genome research, 13(11), 2498–2504.

Sikora, J. and Ouagazzal, A.-M. (2021). Synaptic zinc: an emerging player in parkinson’s disease. International Journal of Molecular Sciences, 22(9), 4724.

Singh, S. S. et al. (2020). Nf-*κ*b-mediated neuroinflammation in parkinson’s disease and potential therapeutic effect of polyphenols. Neurotoxicity Research, 37(3), 491–507.

Smajić, S., et al. (2022). Single-cell sequencing of human midbrain reveals glial activation and a parkinson-specific neuronal state. Brain, 145(3), 964–978.

Tang, X. et al. (2021). Indicator regularized non-negative matrix factorization method-based drug repurposing for covid-19. Frontiers in Immunology, 11, 3824.

Teyssou, E. et al. (2022). The amyotrophic lateral sclerosis m114t pfn1 mutation deregulates alternative autophagy pathways and mitochondrial homeostasis. International Journal of Molecular Sciences, 23(10), 5694.

Thygesen, C. et al. (2018). Characterizing disease-associated changes in post-translational modifications by mass spectrometry. Expert Review of Proteomics, 15(3), 245–258.

Travaglio, M. et al. (2023). Increased cysteine metabolism in pink1 models of parkinson’s disease. Disease Models & Mechanisms, 16(1), dmm049727.

Vavasis, S. A. (2010). On the complexity of nonnegative matrix factorization. SIAM Journal on Optimization, 20(3), 1364–1377.

Welch, J. D. et al. (2019). Single-cell multi-omic integration compares and contrasts features of brain cell identity. Cell, 177(7), 1873–1887.

Wishart, D. S. et al. (2018). Drugbank 5.0: a major update to the drugbank database for 2018. Nucleic Acids Research, 46(D1), D1074–D1082.

Wishart, D. S. et al. (2022). Hmdb 5.0: the human metabolome database for 2022. Nucleic Acids Research, 50(D1), D622–D631.

Yang, J. et al. (2008). Non-negative graph embedding. In 2008 IEEE Conference on Computer Vision and Pattern Recognition, pages 1–8. IEEE.

Yang, W. et al. (2020). Current and projected future economic burden of parkinson’s disease in the us. npj Parkinson’s Disease, 6(1), 1–9.

