## Supplementary Materials for "Multi-omics integration of scRNA-seq time series data predicts new intervention points for Parkinson’s disease"

This Supplementary Materials contains Supplementary Sections 1 to 8, Supplementary Figures 1 to 11 and Supplementary Tables 1 to 6.

### 1 Multiplicative update rules

To obtain the matrix factors of our NetSC-NMTF model described in Section 5.2, we solve the following continuous and non-linear minimization problem:

$$\min_{G_1 \geq 0, G_2 \geq 0, S_i, S_5} F(G_{1,2}, S_{1,..,5}) = \min_{G_1 \geq 0, G_2 \geq 0, S_i, S_5} \sum_{i=1}^{4} \parallel A_i - G_1 S_i G_1^T \parallel_F^2 + \parallel E - G_1 S_5 G_2^T \parallel_F^2 \quad (1)$$

where $\|\|_F$ denotes the Frobenius norm. Furthermore, to ease interpretability, we constrain the embedding matrices $G_1$ and $G_2$ to be non-negative (i.e., the semi-NMTF [5]). To solve our optimization problem, we derive the Karush-Kuhn-Tucker (KKT) conditions, which are necessary for a solution to be optimal [20]:

$$\begin{aligned} \frac{\partial F}{\partial G_1} = & \sum_{i=1}^{4}((-A_i^T G_1 S_i + A_i G_1 S_i^T) + (G_1 S_i G_1^T G_1 S_i^T + G_1 S_i^T G_1^T G_1 S_i)) \\ & - E G_2 S_5^T + G_1 S_5 G_2^T G_2 S_5^T - \eta_1 = 0, \end{aligned} \quad (2)$$

$$\frac{\partial F}{\partial G_2} = -E^T G_1 S_5 + G_2 S_5^T G_1^T G_1 S_5 - \eta_2 = 0, \quad (3)$$

$$\frac{\partial F}{\partial S_1} = -G_1^T A_1 G_1 + G_1^T G_1 S_1 G_1^T G_1 = 0, \quad (4)$$

$$\frac{\partial F}{\partial S_2} = -G_1^T A_2 G_1 + G_1^T G_1 S_2 G_1^T G_1 = 0, \quad (5)$$

$$\frac{\partial F}{\partial S_3} = -G_1^T A_3 G_1 + G_1^T G_1 S_3 G_1^T G_1 = 0, \quad (6)$$

$$\frac{\partial F}{\partial S_4} = -G_1^T A_4 G_1 + G_1^T G_1 S_4 G_1^T G_1 = 0, \quad (7)$$

$$\frac{\partial F}{\partial S_5} = -G_1^T E G_2 + G_1^T G_1 S_5 G_2^T G_2 = 0, \quad (8)$$

$$\eta_1, \eta_2, G_1, G_2 \geq 0, \quad (9)$$

$$\eta_1 \odot G_1 = 0, \quad (10)$$

$$\eta_2 \odot G_2 = 0, \quad (11)$$

where $\odot$ is the Hadamard (element-wise) product, and $\eta_1$ and $\eta_2$, are the dual variables for the primal constraints $G_1 \geq 0$ and $G_2 \geq 0$, respectively. By fixing $G_1$ and $G_2$, we can directly compute each $S_i$, by solving the stationary conditions for $S_i$ derived above, to get the following closed formulas:

$$S_1 = (G_1^T G_1)^{-1} G_1^T A_1 G_1 (G_1^T G_1)^{-1} \tag{12}$$

$$S_2 = (G_1^T G_1)^{-1} G_1^T A_2 G_1 (G_1^T G_1)^{-1} \tag{13}$$

$$S_3 = (G_1^T G_1)^{-1} G_1^T A_3 G_1 (G_1^T G_1)^{-1} \tag{14}$$

$$S_4 = (G_1^T G_1)^{-1} G_1^T A_4 G_1 (G_1^T G_1)^{-1} \tag{15}$$

$$S_5 = (G_1^T G_1)^{-1} G_1^T E G_2 (G_2^T G_2)^{-1} \tag{16}$$

Then, we derive the following multiplicative update rules for $G_1$ and $G_2$ to solve the KKT conditions above [20]:

$$G_{1_{(j,l)}} \leftarrow G_{1_{(j,l)}} \sqrt{\frac{\sum_{i=1}^{4}((A_i^T G_1 S_i)_{j,l}^{+} + G_1(S_i G_1^T G_1 S_i)_{j,l}^{-}) + (E G_2 S_5^T)_{j,l}^{+} + G_1(S_5 G_2^T G_2 S_5^T)_{j,l}^{-}}{\sum_{i=1}^{4}((A_i^T G_1 S_i)_{j,l}^{-} + G_1(S_i G_1^T G_1 S_i)_{j,l}^{+}) + (E G_2 S_5^T)_{j,l}^{-} + G_1(S_5 G_2^T G_2 S_5^T)_{j,l}^{+}}} \tag{17}$$

$$G_{2_{(j,l)}} \leftarrow G_{2_{(j,l)}} \sqrt{\frac{(E^T G_1 S_5)_{j,l}^{+} + G_2(S_5^T G_1^T G_1 S_5)_{j,l}^{-}}{(E^T G_1 S_5)_{j,l}^{-} + G_2(S_5^T G_1^T G_1 S_5)_{j,l}^{+}}}, \tag{18}$$

where $(j, l)$ represents an entry value of $G_1$ or $G_2$. We start from initial solutions, $G_{1_{init}}$ and $G_{2_{init}}$, and iteratively use Equations 12-18 to compute new matrix factors $G_1, G_2$ and $S_{i \in \{1,..,5\}}$ until convergence. To generate initial solutions for $G$ matrices, we apply: 1) random initialization when choosing the optimal number of dimensions of the latent embedding spaces (see Supplementary Section 3.1), or 2) an SVD-based strategy for the final integration of the data, as it makes the solver deterministic while reducing the number of iterations that are needed to achieve convergence [21] (see Section 5.2). We perform the minimization procedure eight times in total, once for each cell condition. The algorithm converges (i.e., the stop criterion (defined in Section 5.2) is met) between 240 and 350 iterations (e.g., Supplementary Figure 1), depending on the cell condition.

### 2 Enrichment analysis

To assess if the integration framework captures biological and functional relations between genes, we apply a clustering algorithm on the resulting gene matrix factors and measure if these clusters are significantly enriched in a set of annotations biological annotations (Gene Ontology (GO) [2], KEGG pathways (KP) [10] and Reactome pathways (RP) [9]).

We compute the probability, $p$, that an annotation is enriched in a cluster of genes by using a hypergeometric test (i.e., sampling without replacement strategy) [23]:

$$p = 1 - \sum_{i=0}^{X-1} \binom{K}{i} \binom{M-K}{N-i} / \binom{M}{N} \tag{19}$$

where $N$ is the number of annotated genes in the cluster, $X$ is the number of genes in the cluster annotated with the given annotation, $M$ is the number of annotated genes in the network, and $K$ is the number of genes in the network annotated with the given annotation. Then, we adjust all *p-values* for multiple hypothesis testing using the Benjamini-Hochberg procedure [3]. All annotations with an adjusted *p-value* $\leq 0.05$ are considered to be statistically significantly enriched. The clustering quality is measured by: 1) the percentage of clusters with at least one enriched annotation, out of all non-empty clusters, and 2) the percentage of genes with at least one of their annotations enriched in their clusters, out of all annotated genes.

### 3 Model parameters

#### 3.1 Choosing the number of dimensions

The number of dimensions ($k_1$ and $k_2$, in our case) that define the latent embedding spaces are key parameters of NMTF-based models. Since there is no universally accepted procedure for obtaining the dimension parameters, we use the procedure inspired by Brunet et al. [4]. The method consists of computing a dispersion coefficient of the gene clusterings obtained for multiple combinations of $k_1$ and $k_2$ and different runs of NMTF using different random initial solutions, and selecting a combination of $k_1$ and $k_2$ where the dispersion coefficient is the highest, indicating cluster stability.

Starting from a random initial solution, we run our NetSC-NMTF algorithm ten times for every cell condition and combination of $k_1$ and $k_2$, where $k_1 \in \{25, 50, 75, 100, 125, 150, 175, 200, 250, 275\}$ and $k_2 \in \{10, 20, 30, 40, 50, 60, 70, 80, 90, 100\}$. We choose the range of $k_1$ based on two underlying principles. The first one comes from the field of network embeddings, where the commonly used sizes for the lower-dimensional embedding spaces are between 100 and 250 [14, 8]. The second one is a widely used heuristic formula (i.e., "rule of thumb"), $k = \sqrt{n/2}$, where $k$ is the number of dimensions and $n$ is the number of objects (genes or single cells) that should be embedded. Therefore, we start from the initial dimension of 25, which is far below the 'rule of thumb" value (around 100, depending on the cell condition) to ensure that the optimal value of $k_1$ is inside the range and not on its border. To embed SCs in lower-dimensional spaces, popular embedding methods such as Seurat [27] and LIGER [29] use dimensions in the range of 20 to 50. To ensure that we find the most optimal value of $k_2$, we explore $k_2$ values by extending this commonly used range from 10 to 100.

For each cell condition, NetSC-NMTF run and ($k_1$, $k_2$) pair, we apply a k-means clustering procedure [18] (with ten runs) to the $G_1$ matrix factor, clustering the genes in $k_1$ number of clusters and to the $G_2$ matrix factor clustering the SCs in $k_2$ number of clusters. We use the obtained gene clusters to generate an $n \times n$ connectivity matrix $C_1$, and the SC clusters to generate an $m \times m$ connectivity matrix $C_2$, where $n$ is the number of genes and $m$ is the number of single cells. Entries in the connectivity matrix $C_1$ are $C_1[u][v] = 1$ if genes $u$ and $v$ belong to the same cluster, or $C_1[u][v] = 0$ otherwise. Similarly, entries in the connectivity matrix $C_2$ are $C_2[u][v] = 1$ if single cells $u$ and $v$ belong to the same cluster, or $C_2[u][v] = 0$ otherwise. Then, for every cell condition and $k_1$ and $k_2$ pair, we individually compute the average of all connectivity matrices $C_1$ and $C_2$ across all runs, $\overline{\mathrm{C_1}}$ and $\overline{\mathrm{C_2}}$, and evaluate the stability of clustering of $G_1$ and $G_2$ matrices with the dispersion coefficient: $\rho_{k_i} = \frac{1}{n^2} \sum_{j=1}^{n} \sum_{l=1}^{n} 4(\overline{\mathrm{C_{i_{[j,l]}}}} - \frac{1}{2})^2$, where $i \in \{1, 2\}$, and $[j, l]$ is an entry of a $\overline{\mathrm{C_1}}$ or a $\overline{\mathrm{C_2}}$ matrix. To choose the best $k_1$ and $k_2$ parameter pair for a cell condition, we calculate the mean of condition-specific dispersion coefficients of $\rho_{k_1}, \rho_{k_2}$ with $mean_{\rho_{k_1},\rho_{k_2}} = \frac{\rho_{k_1}+\rho_{k_2}}{2}$, for all $k_1$ and $k_2$ combinations and identify the one for which $mean_{\rho_{k_1},\rho_{k_2}}$ is at its maximum. The dispersion coefficient is in the [0,1] range, where 0 means that clusters across all runs are different, and 1 means that they are identical.

We find that the $mean_{\rho_{k_1},\rho_{k_2}}$ is maximum, or very close to the maximum, for many combinations of $k_1$ and $k_2$, suggesting that our model is robust to the choice of the dimension parameters (see Supplementary Figures 2 and 3). Additionally, we observe that there is a plateau for the $mean_{\rho_{k_1},\rho_{k_2}}$, so that after increasing $k_1$ and $k_2$, the $mean_{\rho_{k_1},\rho_{k_2}}$ remains stable. To balance between accuracy (clusters are more stable at higher dimensions) and interpretability (clusters contain more biological meaning at lower dimensions), we identify those combinations of $k_1$ and $k_2$ for each cell condition where we observe a plateau for the $mean_{\rho_{k_1},\rho_{k_2}}$ (see Supplementary Figures 2 and 3). This is achieved for $k_1$ between 75 and 125, and $k_2$ between 40 and 60, with $mean_{\rho_{k_1},\rho_{k_2}}$ ranging between 0.905 and 0.932. For the combination of $k_1$ and $k_2$ used for each cell condition, see Supplementary Table 3.

#### 3.2 Integrating single-cell expression data with molecular networks captures the functional organization of cell conditions

To validate our NetSC-NMTF data-integration model (that integrates a single cell expression matrix, $E$, together with molecular interaction networks), we assess how well it captures the functional organization of cell conditions and investigate if all molecular networks contribute to the decomposition. To this aim, we perform the following gene clustering and functional enrichment analysis. We obtain clusters of genes for each cell condition by applying k-means clustering [18] to the corresponding condition-specific $G_1$ matrix (i.e., a set of gene embedding vectors). We perform the clustering step for 16 combinations of SC expression data and molecular networks (i.e., integration scenarios), where the $E$ matrix is integrated alone or with one or multiple molecular networks. We apply the k-means algorithm ten times on the $G_1$ matrices obtained from all integration scenarios and for every cell condition to account for the non-deterministic behaviour of the k-means algorithm. Then, we perform an enrichment analysis (described in Supplementary Section 2) of the resulting clusters in *DisGeNet PD genes*. We choose the run for each integration scenario and cell condition where the significantly enriched clusters contain the highest number of *DisGeNet PD genes*. We hypothesize that the more *DisGeNet PD genes* a cluster contains, the higher the probability is that other genes in that cluster are also PD-relevant. We use this hypothesis to validate the biological relevance of gene clusters by first taking the k-means clustering run that is most enriched in PD genes and then evaluating if those most PD-related clusters are also biologically relevant. We use this notion in the 2-step downstream method to cluster genes based on their relationship with PD and then investigate those clusters to find new PD-relevant biology. Hence, our biological validation pipeline follows this hypothesis by first taking the k-means clustering run that is most enriched in PD genes and then evaluating if those most PD-related clusters are also biologically relevant.

To measure the quality of gene clustering and evaluate their biological meaning, we perform the enrichment analysis in GO, KP and RP biological annotations (described in Supplementary Section 2) of the k-means clusters from the best run for each integration scenario and cell condition. More precisely, we compute the percentage of enriched clusters (see Supplementary Figure 4, a) and the percentage of genes (see Supplementary Figure 4, b) with enriched GO, KP and RP biological annotations. High enrichments in both measurements indicate that clusters show relevant biological functionality.

For many integration scenarios, we observe the following trends across all cell conditions:

1. KP annotations - percentages of enriched clusters are around 80%, and percentages of genes with enriched annotations are 35%;

2. RP annotations - percentages of enriched clusters are around 90%, and percentages of genes with enriched annotations are around 45%;

3. GO annotations - percentages of enriched clusters go as high as 100% (all clusters are enriched), and percentages of genes with enriched annotations are between 40% and 50%.

These results indicate that five integration scenarios across all cell conditions (E+P+C, E+C+M, E+G+C+M, E+P+C+M, E+P+M+C+G; E is an expression matrix, C is a COEX network, P is a PPI network, M is a MI network, and G is a GI network) achieve the best and comparably high cluster and gene enrichments, that vary by a small percentage ($< 2\%$) across the five different integration scenarios. In particular, we observe that the clusters of genes coming from integrating all input data (E+P+M+C+G) are statistically significantly enriched in KPs, RPs, and GO, with an average amount of enriched clusters of 83.3%, 94% and 100%, and of enriched genes of 39.8%, 45.6% and 53.2%, respectively. This scenario leads to functionally coherent clusters across cell conditions and demonstrates the utility of integrating expression data with all available molecular networks. The results also show that decomposing the expression matrix alone (E scenario in Supplementary Figure 4) does not yield functionally coherent spaces (percentages of enriched clusters and genes with enriched annotations are less than 5%), further demonstrating the power of our data fusion model.

Our results show that leveraging the information content of all molecular networks with SC expression data leads to one the most coherent representations of cell functioning where genes embedded close to each other are functionally related - further highlighting the value of data fusion [15, 7]. Therefore, we choose the integration scenario based on the integration of all data so that the embeddings are informed by all the available data. Thus, we produce gene embeddings of each cell condition by integrating all condition-specific molecular networks with scRNA-seq data.

### 4 "Movement" of ***DisGeNet PD genes*** projected in the SC embedding spaces

In our study, we hypothesize that the functional organization of *DisGeNet PD genes* is more altered than that of background genes between time point-matching PD and control cell conditions. In our methodology, this means that *DisGeNet PD genes* should have greater changes in their relative positions between the two cell conditions ("movement"; see Section 5.3) than non-*DisGeNet PD genes* (i.e., background genes) at a specific time point.

In this section, we assess if this "movement" property of *DisGeNet PD genes* is better captured by directly using the "gene embeddings" from $G_1$ or by using the "gene embeddings" from $U = G_1 * S_5$. Matrix $U$ is the projection of the "gene embeddings" in the SC embedding space spanned by $G_2$ (as described in Section 5.3). We measure this by calculating the distribution of the "movement" of *DisGeNet PD genes* and of background genes for each time point and type of "gene embedding".

Next, we compare the two time point-matching distributions across all time points and for each type of "gene embedding" using a one-sided Mann-Whitney U (MWU) test (with a significance level of 0.05). We observe that the "movement" of *DisGeNet PD genes* based on "gene embeddings" of $G_1$ are only significantly higher for day 21 ($p$-$value = 1.04e^{-07}$), unlike their "movement" based on "gene embeddings" of $U$, where *DisGeNet PD genes* have higher "movement" across all time points ($p$-$value \leq 1.65e^{-5}$).

The results suggest that projecting the raw "gene embeddings" in the SC embedding space emphasizes the contribution of a biological condition of SCs, making "gene embeddings" of stage-specific PD and control cell conditions more distinct and easier to compare. Therefore, since $U$ matrices better capture the functional organization of *DisGeNet PD genes*, we use them to track the "movement" of genes.

#### 5 Stage-specific PD predictions

First, we exploit the properties of *DisGeNet PD genes* to define ***Stage-specific PD predictions***, which we then validate in the literature. For this purpose, we apply the first step of our 2-step downstream methodology (Section 5.4) to analyze the clusters of genes that are significantly enriched in *DisGeNet PD genes* (Section 2.1) and generate four sets of ***Stage-specific PD predictions*** that contain the following number of genes: $\text{PD}_{\text{D0}}$= 1,333, $\text{PD}_{\text{D6}}$ = 1,268, $\text{PD}_{\text{D15}}$ = 1,281 and $\text{PD}_{\text{D21}}$ = 1,017.

To assess if the ***Stage-specific PD predictions*** can be used to identify novel PD-associated genes, we verify that they follow the same "movement" property as *DisGeNet PD genes*, i.e., they have larger "movement" than background genes (Section 2.1). Thus, for each time point, we compute the distributions of "movement" of ***Stage-specific PD predictions*** and background genes using the methodology described in Section 5.3. By comparing the two distributions (a one-sided MWU test with a 0.05 significance threshold), we observe a significantly higher "movement" of ***Stage-specific PD predictions*** across all time points (Supplementary Figure 8; $p$-$value \leq 4.73e^{-10}$), same as the "movement" of *DisGeNet PD genes*.

We assess the PD relevance of our ***Stage-specific PD predictions*** by checking if they have been significantly associated with PD in the literature. For this purpose, we use an automated PubMed publications search to count the co-occurrence of each gene in a prediction set with the term “Parkinson’s disease” in PubMed publications (see Section 5.5). Using a MWU test, we compare the distribution of the literature co-occurrence of each set of ***Stage-specific PD predictions*** with its corresponding background. We observe that all four sets of ***Stage-specific PD predictions*** are significantly more associated with PD than the background genes ($p\text{-}value \leq 9.92e^{-21}$, Supplementary Figure 9), confirming the relevance of our results for PD. Additionally, we observe that all sets of ***Stage-specific PD predictions*** are significantly enriched in the PD-related genes, with $p\text{-}value \leq 2.38^{-08}$ (see Supplementary Figure 11).

Taken together, we demonstrate that our ***Stage-specific PD predictions*** associated with all four time points of cell development are related to PD.

### 6 Sampling with replacement

To see if our 2-step downstream method (Section 5.4) produces more ***Core PD predictions*** than could be obtained by intersecting random sets of genes, we apply a sampling with replacement technique. First, From the genes that are expressed in a PD and control cell conditions at a given time-point, we randomly sample 4 sets of genes having the same sizes as the corresponding sets of ***Stage-specific PD predictions***. Then, we intersect those random sets of predictions to see how many times their overlap is larger than or equal to our***Core PD predictions*** (success) by calculating the $p\text{-}value = \frac{s+1}{r+1}$, where $s$ is the number of successes and $r$ the number of repetitions. If $p\text{-}value \leq 0.05$, we observe that the overlap between ***Stage-specific PD predictions*** is statistically significantly higher than that of random sets of genes, showing that such a consensus approach is possible and represents a promising strategy for discovering new PD-associated genes.

### 7 Comparison with other methods

To demonstrate that our 2-step downstream methodology produces gene predictions that are more relevant to PD than those obtained by other algorithms, we compare our methodology with LIGER

[29], which uses an integrative NMF approach to identify shared and specific sources of variation across datasets. To show the benefit of our 2-step downstream methodology that uses both the clustering and the movement properties of *DisGeNet PD genes* (defined in Section 5.1.2), we compare its ability to predict PD-associated genes to those of more straightforward methods that use these properties individually.

#### 7.1 Our framework vs LIGER

To obtain stage-specific sets of predictions using the LIGER method (an NMF-based approach), we follow the tutorial "Integrating Multiple Single-Cell RNA-seq Datasets" [28], which allows us to identify gene markers based on differential expression analysis between a PD and control cell condition. We compare our approach to LIGER, because it is most closely related to our framework. An important difference with our method is that it is not designed to integrate scRNA-seq data with molecular networks, which is why we apply LIGER to each PD and control pair of expression matrices. We use LIGER with its recommended settings from [29], but with 100 as the dimension of factorization parameter. We choose the dimension of 100 to produce embedding spaces of genes with dimensions similar to what we obtain with our NetSC-NMTF model (from 75 to 125 dimensions, Supplementary Section 3). For time points at day 0, 6, 15 and 21, we obtain sets with 2,575, 1,602, 1,986 and 1,500 genes, respectively. To determine the PD-relevance of each prediction set, we use a MWU test to compare the distribution of the co-occurrence of its genes with the term "Parkinson's disease", in the Pubmed publications, with: 1) the distribution of the co-occurrence of the background genes expressed at a particular time point, and 2) the distribution of the co-occurrence of the ***Stage-specific PD predictions*** expressed at a particular time point (Section 5.5). We find that all four sets of gene predictions obtained by LIGER are more cited in literature with the term "Parkinson's disease" (*p-value* $\leq 6.43e^{-9}$) than the corresponding background genes. However, we observe that no set of LIGER predictions co-occurs more than the ***Stage-specific PD predictions***. Then, we enrich each set individually in the PD-related genes (defined in Section 5.5) and observe that predictions corresponding to time points at day 0, 6 and 21 are significantly enriched (*p-value* $\leq 1.01e^{-4}$), in contrast to all four of our ***Stage-specific PD prediction*** sets.

Finally, we intersect all stage-specific sets to obtain a set equivalent to our ***Core PD predictions***, resulting in a set of 44 gene predictions which is statistically significant (confirmed by doing a sampling with replacement; see Supplementary Section 6), but noticeably lower than our 193 ***Core PD predictions***. We apply the literature co-occurrence validation procedure to this final set of predictions and perform the enrichment analysis in PD-related genes. Neither experiment leads to a significant *p-value* (less than 0.05), thereby failing to confirm the significance of these genes to PD and showing that LIGER cannot be used to uncover novel PD genes relevant across all time points.

#### 7.2 Our framework vs predicting new PD-associated genes based on the property of ***DisGeNet PD genes*** to group together

To predict novel PD-associated genes using the property of *DisGeNet PD genes* to group together, we determine the distance of each non-*DisGeNet PD gene* to each *DisGeNet PD genes* by calculating the Euclidean distance between their gene embedding vectors from the $G_1$ matrix for each PD cell condition. As with ***Stage-specific PD predictions***, we only include non-*DisGeNet PD gene* that are both in PD and control time point-matching cell conditions. We rank the non-*DisGeNet PD genes* according to the smallest distance to any *DisGeNet PD gene*.

For choosing the prediction sets equivalent to ***Stage-specific PD prediction***, we focus on three different cases by taking: 1) the top 5% highest-ranked genes, 2) the top 10% highest-ranked genes, and 3) the same number of the highest-ranked genes as the original ***Stage-specific PD prediction*** sets at matching time points. This results in three groups, each consisting of four stage-specific prediction sets. To see if they are statistically connected to PD, we validate our predictions in the literature by applying the literature co-occurrence validation procedure and performing the enrichment analysis in PD-related genes (Section 5.5). Both experiments show that no set of these predictions is significantly validated in literature because the computed *p-values* are never smaller than or equal to the 0.05 threshold. Because these predictions are not statistically significantly relevant to PD, we do not continue with the downstream pipeline to obtain a set of genes equivalent to our ***Core PD predictions***. Therefore, we conclude that relying only on the first property of *DisGeNet PD genes* does not lead to predictions that are more PD-relevant than those presented in the main paper.

##### 7.3 Our framework vs predicting new PD-associated genes based on the higher "movement" property of *DisGeNet PD genes*

To predict new PD-related genes relying only on the property that *DisGeNet PD genes* move more between PD and control cell conditions than background, we calculate the "movement" between the non-*DisGeNet PD genes* expressed in PD and control cell conditions at a matching time point, ranking them according to the largest "movement". We repeat this for all four time points. For choosing the stage-specific sets of predictions equivalent to the ***Stage-specific PD prediction***, we focus on three different cases by taking: 1) the top 5% highest-ranked genes, 2) the top 10% highest-ranked genes, and 3) the same number of the highest-ranked genes as the original ***Stage-specific PD prediction*** sets at matching time points (as explained in the previous paragraph). We repeat the literature validation analyses described in Section 5.5 and find that all stage-specific sets of predictions (from all three cases) co-occur more with the term "Parkinson's disease" in Pubmed publications than background (Supplementary Table 6), and they are significantly enriched in the PD-related genes (Supplementary Table 6). However, by applying a one-sided MWU test to investigate if the stage-specific sets of predictions co-occur more with the term "Parkinson's disease" in PubMed than the original ***Stage-specific PD predictions***, we observe that no set of these predictions (from any of the three cases) co-occurs more than the ***Stage-specific PD predictions***.

For each case, we similarly obtain the ***Core PD predictions*** by intersecting the stage-specific sets of predictions and rank them according to the average "movement" across all time points, largest first, thereby obtaining sets of 6, 24 and 66 genes, corresponding to the intersection between the top 5% highest-ranked genes, the top 10% highest-ranked genes, and the same number of the highest-ranked genes as the original ***Stage-specific PD prediction*** sets at matching time points, respectively. The number of genes is significantly lower than the 193 ***Core PD predictions*** that we obtain with our 2-step methodology. We validate the predictions in literature (Section 5.5) and find that the predictions of all cases co-occur more with the term “Parkinson’s disease” in Pubmed publications than background genes (Supplementary Table 6). However, by applying a one-sided MWU test to investigate if the sets of gene predictions in the different intersections co-occur more with the term “Parkinson’s disease” in PubMed than the ***Core PD predictions***, we observe that no set of new predictions (from any of the three cases) co-occurs more than the ***Core PD predictions***. The predictions of all three cases are also statistically significantly enriched in the PD-related genes (Supplementary Table 6).

After globally validating all three cases in the literature, we perform the enrichment analysis in KEGG pathways and find that only the predictions in the intersection corresponding to case 3 (i.e., the case where the stage-specific sets of predictions consist of the same number of the highest-ranked genes as the ***Stage-specific PD prediction*** sets at matching time points) are significantly enriched in 10 KEGG pathways. Most importantly, the 66 gene predictions are enriched in the *Parkinson’s disease* pathway and several other neurodegenerative diseases, including *Alzheimer’s disease*, *Huntington’s disease* and *Amyotrophic lateral sclerosis*. To determine if this new set of predictions is more important for the metabolism of PD (metabolism is altered in PD [1]) than our 193 ***Core PD predictions***, we inspect their subgraphs of the general MI network obtained from KEGG. We find that only 2 of the new 66 predictions are present in the MI network, in contrast to the 31 genes of the ***Core PD predictions*** that form a connected component of 29 genes, showing that ***Core PD predictions*** are more involved in metabolic processes than the new set of predictions. Therefore, ***Core PD predictions*** generated by our 2-step downstream methodology are more likely to contain PD-associated genes that participate in PD metabolism. Thereby, we demonstrate that relying only on the second property of *DisGeNet PD genes* leads to fewer predictions that are less PD-relevant than those presented in the main paper.

Overall, in this section, we show that our 2-step downstream method leads to more novel PD-associated gene predictions that are more relevant for PD than the predictions obtained with: 1) LIGER [29], 2) a method based on the distance of non-*DisGeNet PD genes* to *DisGeNet PD genes* in the embedding spaces of PD cell conditions, or 3) an approach based on calculating “movement” of genes between control and PD stage-specific cell conditions.

### 8 Enriched KEGG pathways shared between *DisGeNet PD genes* and *Core PD predictions* are relevant for PD

To further demonstrate that our ***Core PD predictions*** are associated with PD, we examine if the 14 significantly enriched KEGG pathways that are enriched for both *DisGeNet PD genes* and ***Core PD predictions*** are relevant for PD.

*Protein processing in ER*, *carbon metabolism* and *non-alcoholic fatty liver disease* and *Parkinson disease* pathways have already been discussed in Section 2.4 As *central carbon metabolism in cancer* pathway is a subset of the overarching carbon metabolism we argue that it is also relevant for PD. While *glucagon signaling* pathway has not been directly implicated in PD, it contributes to the carbon metabolism and mitochondrial energy production mechanisms, both of which are altered in PD [1, 19]. Therefore, the role of *glucagon signaling* pathway in PD should be investigated in more detail, as it could represent a promising point of intervention for treatment. *The HIF-1 signaling* pathway has been associated to several molecular pathways disrupted in PD such as mitochondrial dysfunction, oxidative stress and protein degradation impairment [12]. The enrichment in the pathways of other neurological diseases (*Huntington disease*, *Alzheimer disease*, *amyotrophic lateral sclerosis* and *prion disease*) is not surprising given that their pathogenesis is characterized by the disruption of many molecular pathways shared with PD, such as the activation of the RAGE receptor, increased endoplasmic reticulum stress and unfolded protein response [22, 6]. Our predictions are also enriched in the pathways of four diseases: *non-alcoholic fatty liver disease*, *diabetic cardiomyopathy*, *Salmonella infection* and *pathogenic Escherichia coli infection.* Some evidence suggests that *Diabetic cardiomyopathy* is also associated with the increased risk of PD [24]; however, more studies would be necessary to elucidate the exact interplay between these two diseases. *Infection with Salmonella* is implicated in a cascade of events that ultimately lead to progressive loss of dopaminergic neurons, a main characteristic of PD [11]. Based on epidemiologic evidence and pathophysiological insights, bacterial infections such as *Escherichia coli* may increase the risk of developing PD [25]. The gut microbiome is actively being investigated for its involvement in PD, as studies show differences in the gut microbiome between healthy individuals and PD patients.

Moreover, more studies suggest that PD may even start in the gut. For example, *Escherichia coli* produces endotoxins that have a reported role in aggregating synuclein and generating toxic synuclein products that can contribute to the development of PD [25]. Further studies are necessary to elucidate the relationship between PD and the gut microbiome, as it could represent a new point of intervention to slow down the disease progression and even target its cause.

The 14 significantly enriched KEGG pathways shared between ***Core PD predictions*** and *DisGeNet PD genes* also provide strong evidence that our set of predictions participates in mechanisms whose disruption promotes PD development. Additionally, further studying the PD importance of the above-mentioned mechanisms could lead to novel treatment strategies for this disease.

### Supplementary Figures

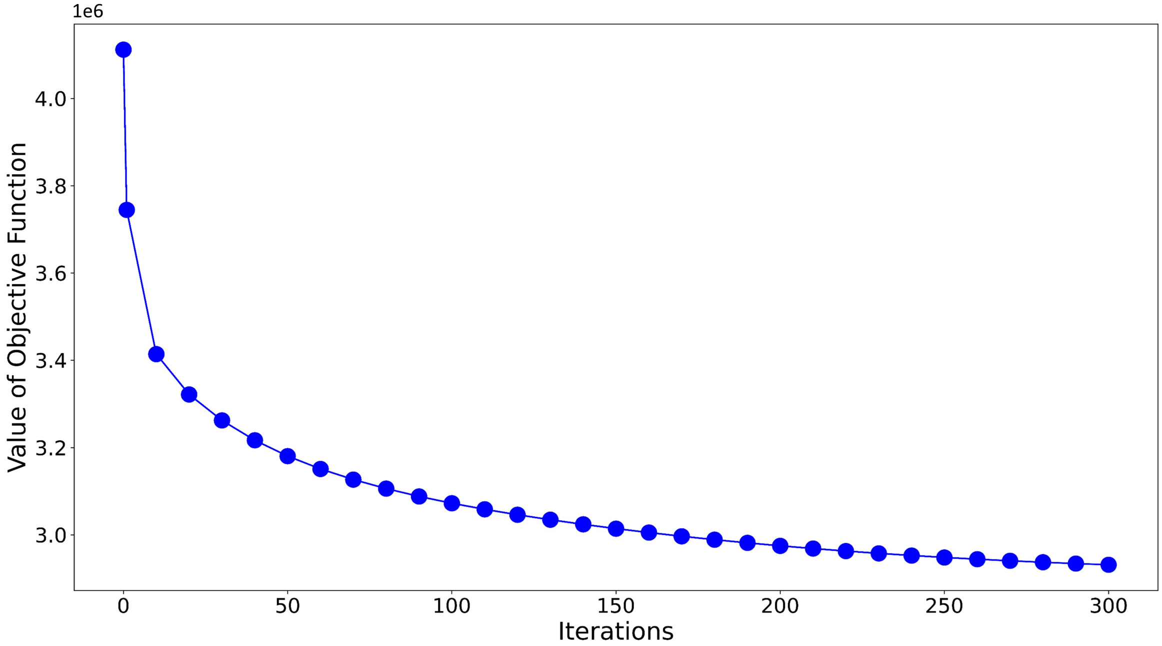


Supplementary Figure 1: **Convergence of the objective function of NetSC-NMTF when integrating PPI, GI, COEX and MI networks with SC expression data of Control$_{\mathbf{D6}}$.**

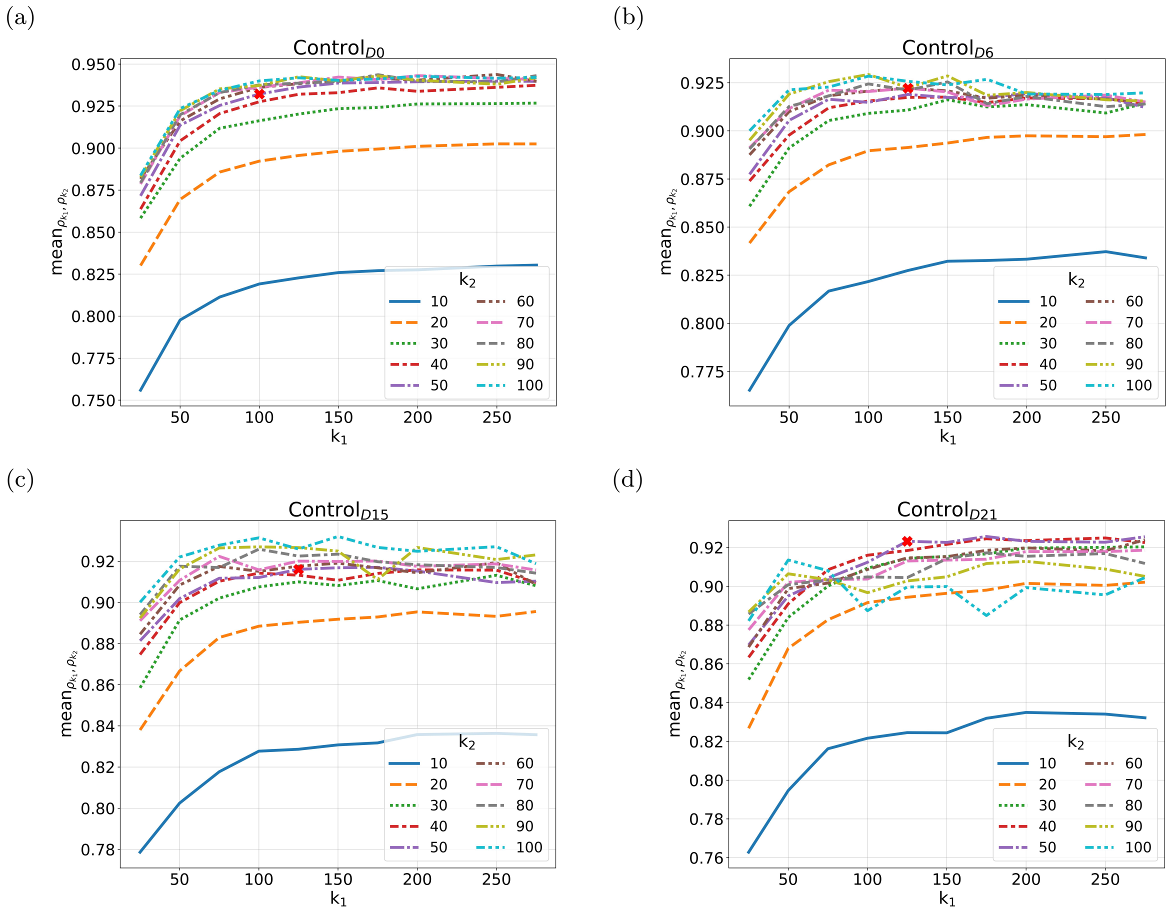


Supplementary Figure 2: **Illustration of grid search results for choosing the values of the dimensions of the embeddings, $k_1$ and $k_2$, based on computing dispersion coefficient for control cell conditions.** The lines represent the mean of the dispersion coefficients across different runs ($mean_{\rho_{k_1},\rho_{k_2}}$) for different values of $k_1$ and $k_2$ for control cell conditions at individual time points (a) $\mathrm{Control_{D0}}$; (b) $\mathrm{Control_{D6}}$; (c) $\mathrm{Control_{D15}}$; (d) $\mathrm{Control_{D21}}$). The red x represents the optimal mean dispersion coefficient with the corresponding $k_1$ and $k_2$ for each control cell condition.

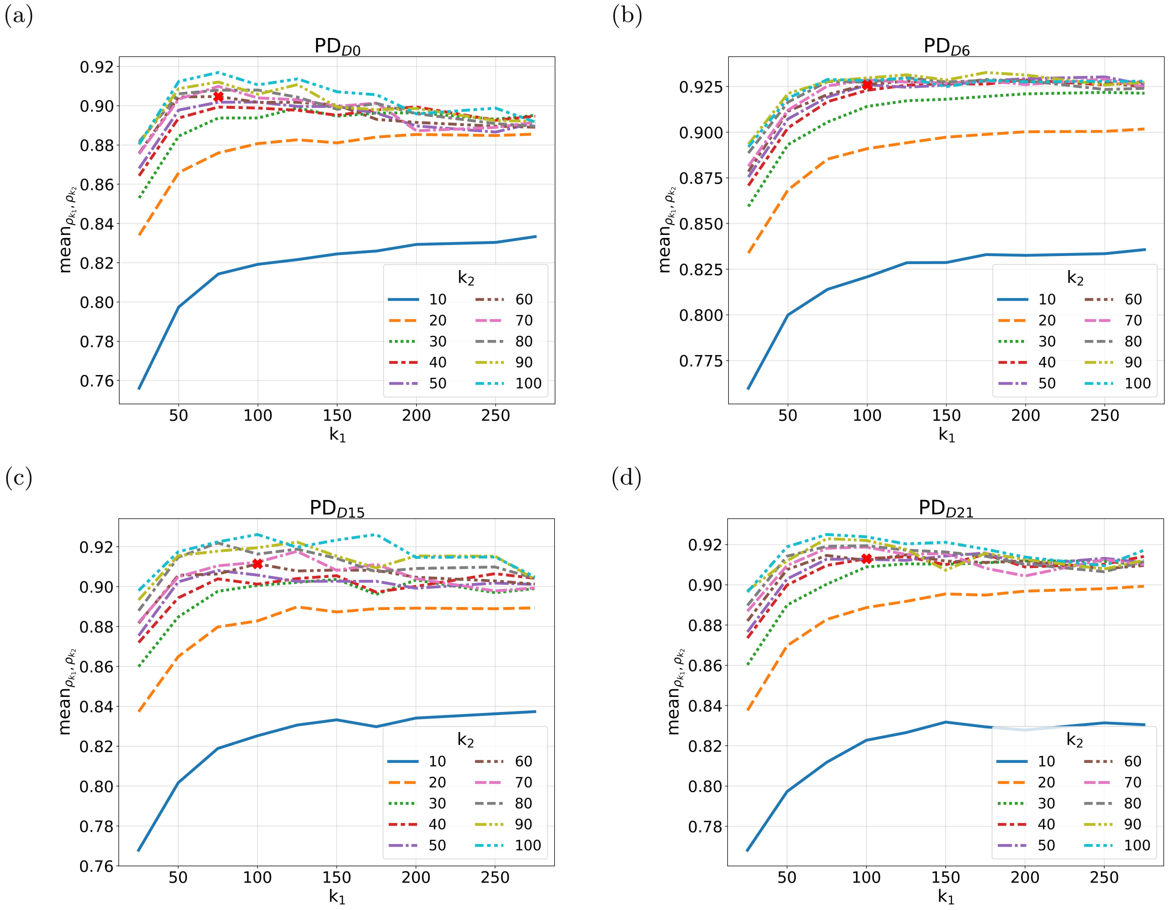


Supplementary Figure 3: **Illustration of grid search results for choosing the values of the dimensions of the embeddings, $k_1$ and $k_2$, based on computing dispersion coefficient for PD cell conditions.** The lines represent the mean of the dispersion coefficients across different runs ($mean_{\rho_{k_1},\rho_{k_2}}$) for different values of $k_1$ and $k_2$ for PD cell conditions at individual time points (a) $\mathrm{PD_{D0}}$; (b) $\mathrm{PD_{D6}}$; (c) $\mathrm{PD_{D15}}$; (d) $\mathrm{PD_{D21}}$). The red x represents the optimal mean dispersion coefficient with the corresponding $k_1$ and $k_2$ for each PD cell condition.

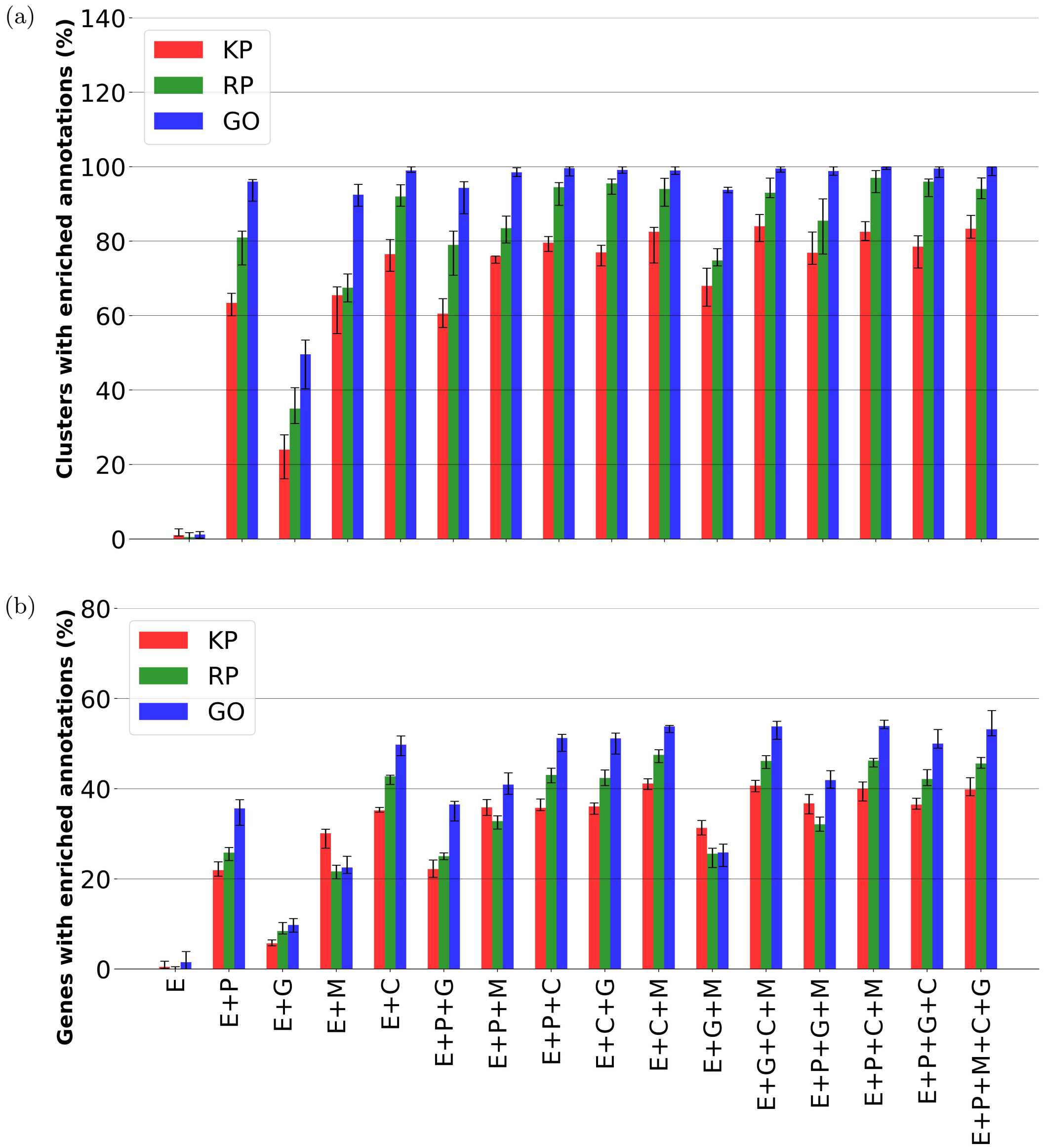


Supplementary Figure 4: **Percentages of (a) enriched clusters and (b) genes with enriched annotations across all cell conditions, calculated using the matrix $G_1$.** We create clusters of genes for 16 combinations of input matrices and investigate their biological functionality to determine if our integration framework produces biologically relevant gene embeddings. **(a)** For each clustering, the bars show the percentage of clusters with at least one enriched annotation in a cluster, out of all non-empty clusters. **(b)** For each clustering, the bars show the percentage of genes with at least one of their annotations enriched in their clusters, out of all annotated genes. Annotations are KEGG pathways (KP), Reactome pathways (RP), and gene ontology terms (GO). The error bars represent the $16^{th}$ and $84^{th}$ percentiles (i.e., percentiles equivalent to the one standard deviation for a normal distribution) of enrichment values, across all cell conditions. E: Expression Matrix; P: protein-protein interaction network; G: genetic interaction network; C: gene co-expression interaction network; M: metabolic interaction network.

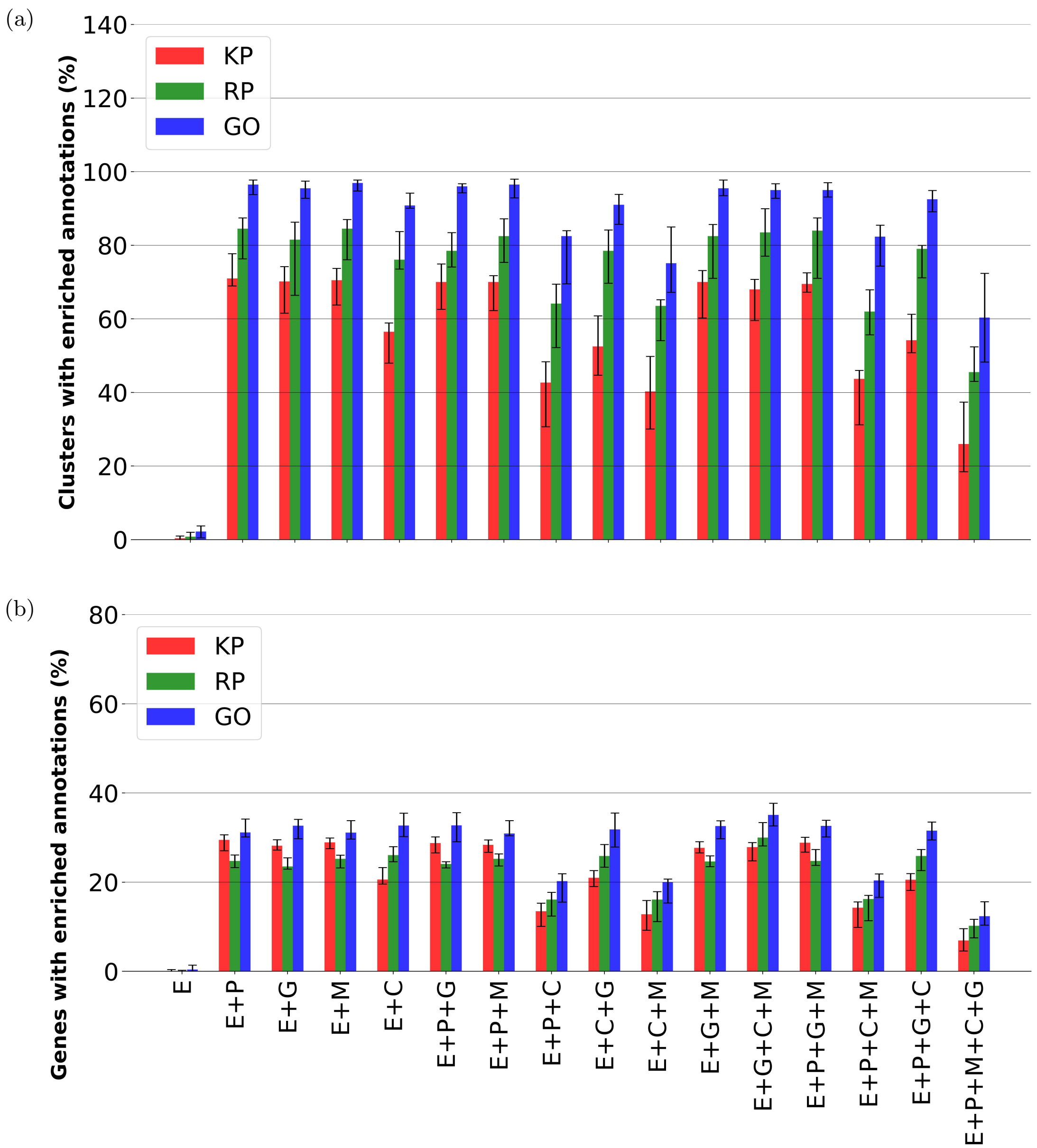


Supplementary Figure 5: **Percentages of (a) clusters and (b) genes with enriched annotations across all cell conditions, calculated using the matrix $U$.** We create clusters of genes for 16 combinations of input matrices and investigate their biological functionality. **(a)** For each clustering, the bars show the percentage of clusters with at least one enriched annotation in a cluster, out of all non-empty clusters. **(b)** For each clustering, the bars show the percentage of genes with at least one of their annotations enriched in their clusters, out of all annotated genes. Annotations are KEGG pathways (KP), Reactome pathways (RP), and gene ontology terms (GO). The error bars represent the $16^{th}$ and $84^{th}$ percentiles (i.e., percentiles equivalent to the one standard deviation for a normal distribution) of enrichment values, across all cell conditions. E: Expression Matrix; P: protein-protein interaction network; G: genetic interaction network; C: gene co-expression interaction network; M: metabolic interaction network.

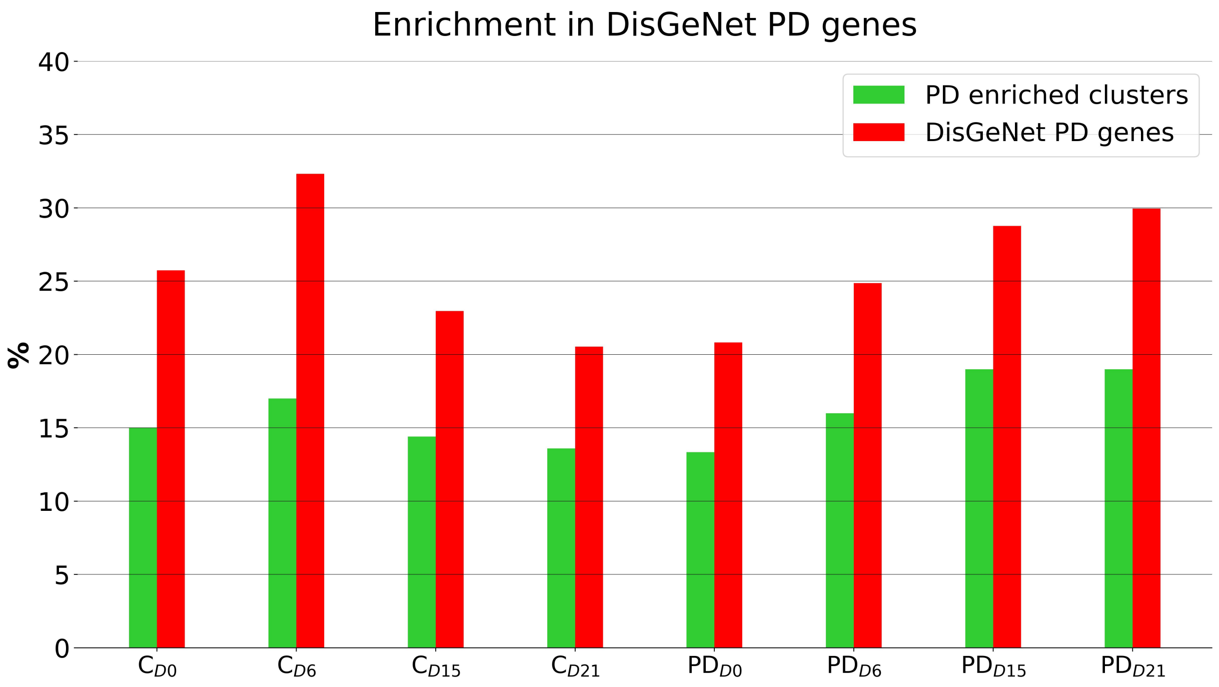


Supplementary Figure 6: **Percentage of clusters significantly enriched in *DisGeNet PD genes* (green) and the percentage of *DisGeNet PD genes* they contain (red) for each individual cell condition.** To obtain the clusters, we apply k-means clustering to the gene embeddings of NetSC-NMTF of each cell condition. C: Control; PD: Parkinson's disease.

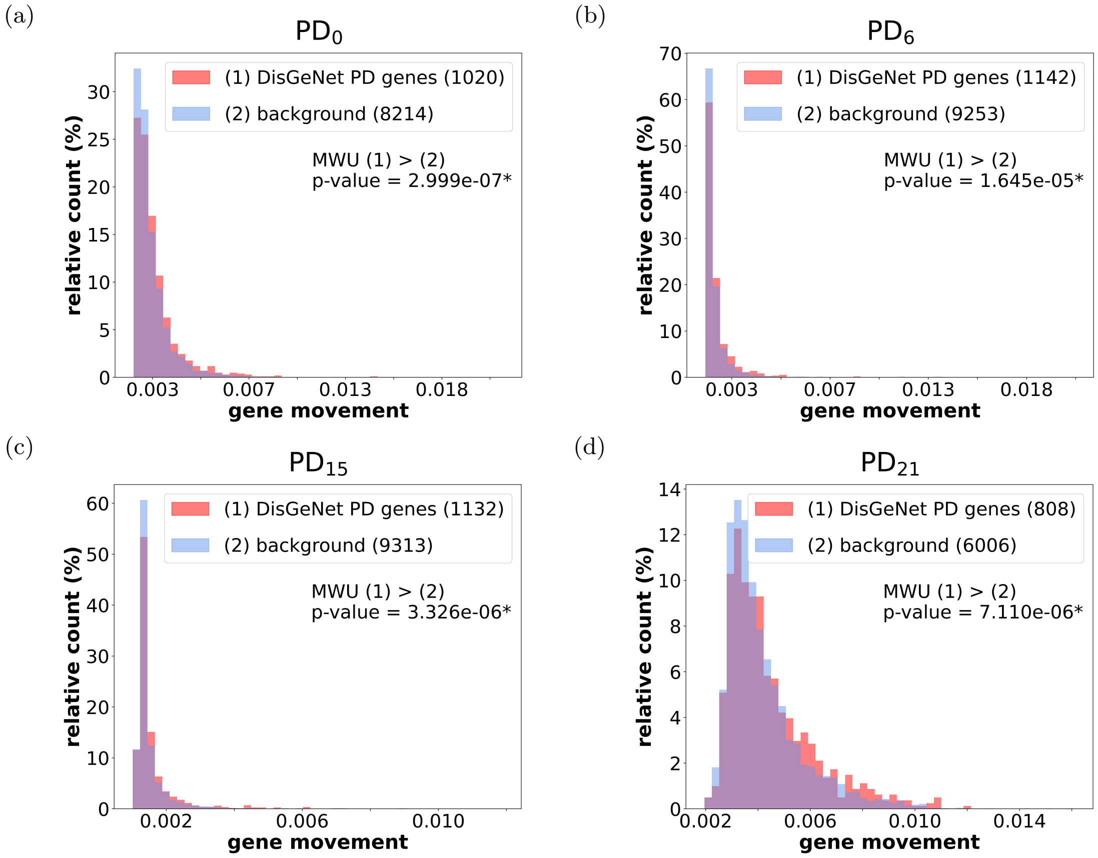


Supplementary Figure 7: **Comparing the "movement" of *DisGeNet PD genes* with the background genes expressed at a particular time point.** **(a)** For each gene expressed in both Control and PD cell conditions at day **(a)** 0, **(b)** 6, **(c)** 15, or **(d)** 21, we calculate the "movement" between the gene's embedding vectors of the two $G_1$ matrices. A gene can either belong to a set of *DisGeNet PD genes* (red) or non-*DisGeNet PD genes* (*background*) (blue) that are expressed at a time point in both PD and control cell conditions. The graph shows normalized "movement" distribution histograms of the two sets of genes, where each bin of the distribution histogram is normalized by dividing it by the number of all values in the distribution. We perform a one-sided Mann-Whitney U test (MWU) (with a significance level of 0.05) to test whether the "movement" distribution of *DisGeNet PD genes* is significantly larger than for *background* (with *p-value* $< 0.05$ indicated by *)

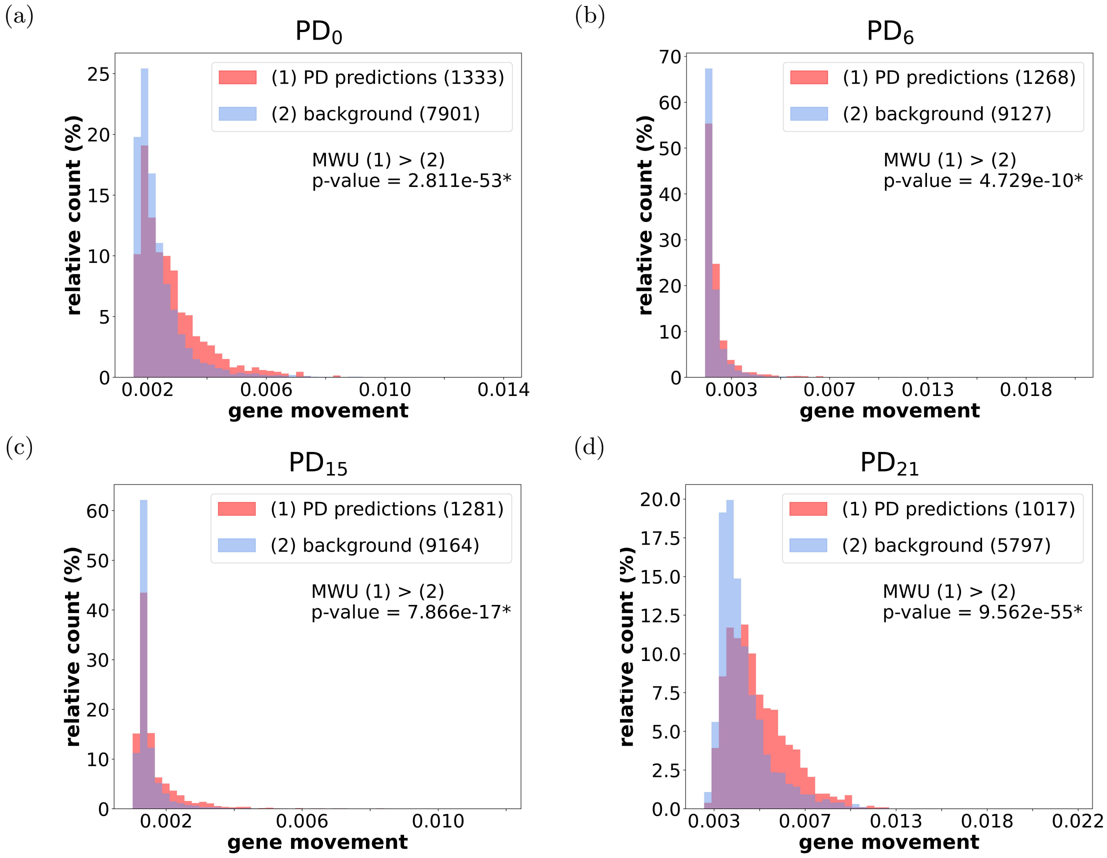


Supplementary Figure 8: **Comparing "movement" of *Stage-specific PD predictions* with the background genes expressed at a particular time point.** For each gene expressed in both Control and PD cell conditions at day **(a)** 0, **(b)** 6, **(c)** 15, or **(d)** 21, we calculate the "movement" between the gene's embedding vectors of the two $G_1$ matrices. A gene can either belong to a set of ***Stage-specific PD predictions*** (red) or *background* (the rest of the non-*DisGeNet PD genes* that are expressed at a time point in both PD and control cell condition) (blue). The graph shows normalized "movement" distribution histograms of the two sets of genes, where each bin of the distribution histogram is normalized by dividing it by the number of all values in the distribution. We perform a one-sided Mann-Whitney U test (MWU) (with a significance level of 0.05) to see if the "movement" distribution of ***Stage-specific PD predictions*** is larger than for *background* (with *p-value* $< 0.05$ indicated by *).

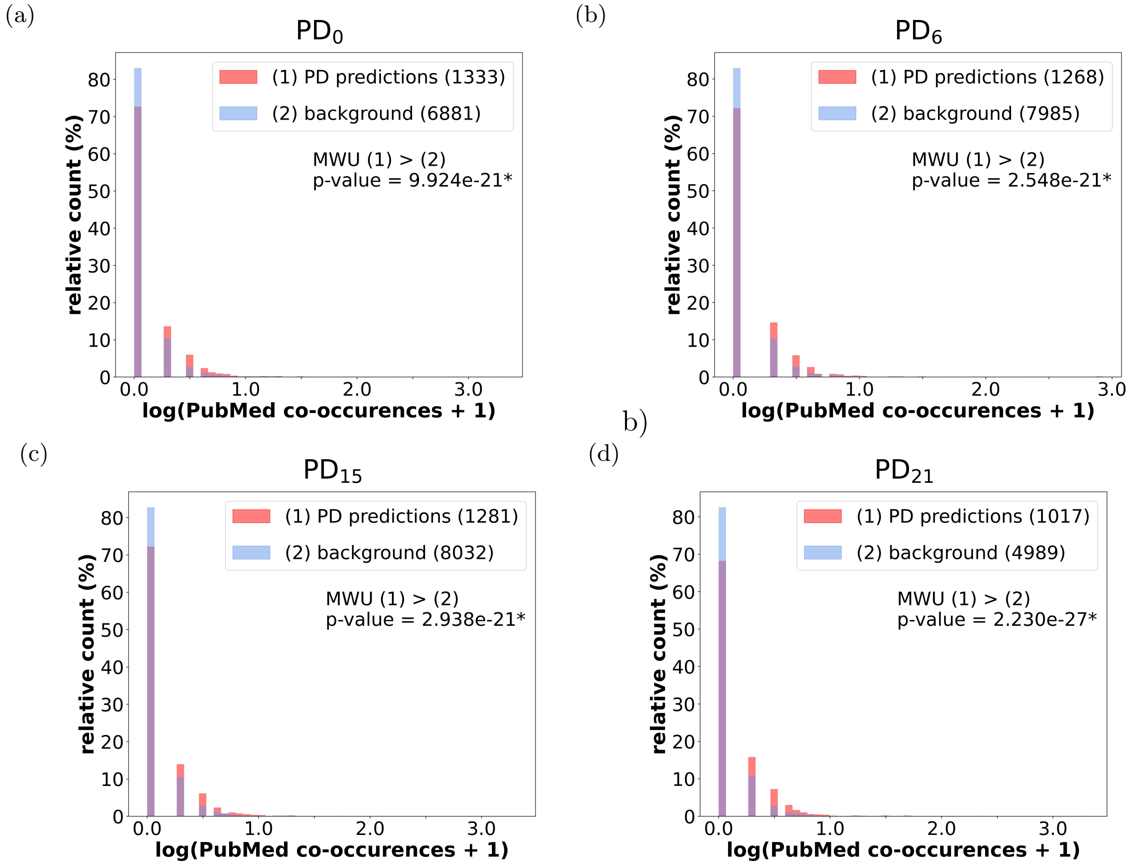


Supplementary Figure 9: **The log-transformed and normalized co-occurrence distributions of *Stage-specific PD predictions* with the term "Parkinson's disease" in PubMed compared to *background* expressed at a particular time point.** We determine the co-occurrence of each gene from a set of ***Stage-specific PD predictions*** and for each gene from the *background* (genes that are not *DisGeNet PD genes* or ***Stage-specific PD predictions***, which are expressed at a time point in both PD and control cell condition) at day **(a)**0; **(b)** 6; **(c)** 15; and **(d)** 21. A one-sided Mann-Whitney U (MWU) test (with a significance level of 0.05) checks if the co-occurrence distribution of a set of ***Stage-specific PD predictions*** is significantly larger than the *background*(with *p-value* $< 0.05$ indicated by *)

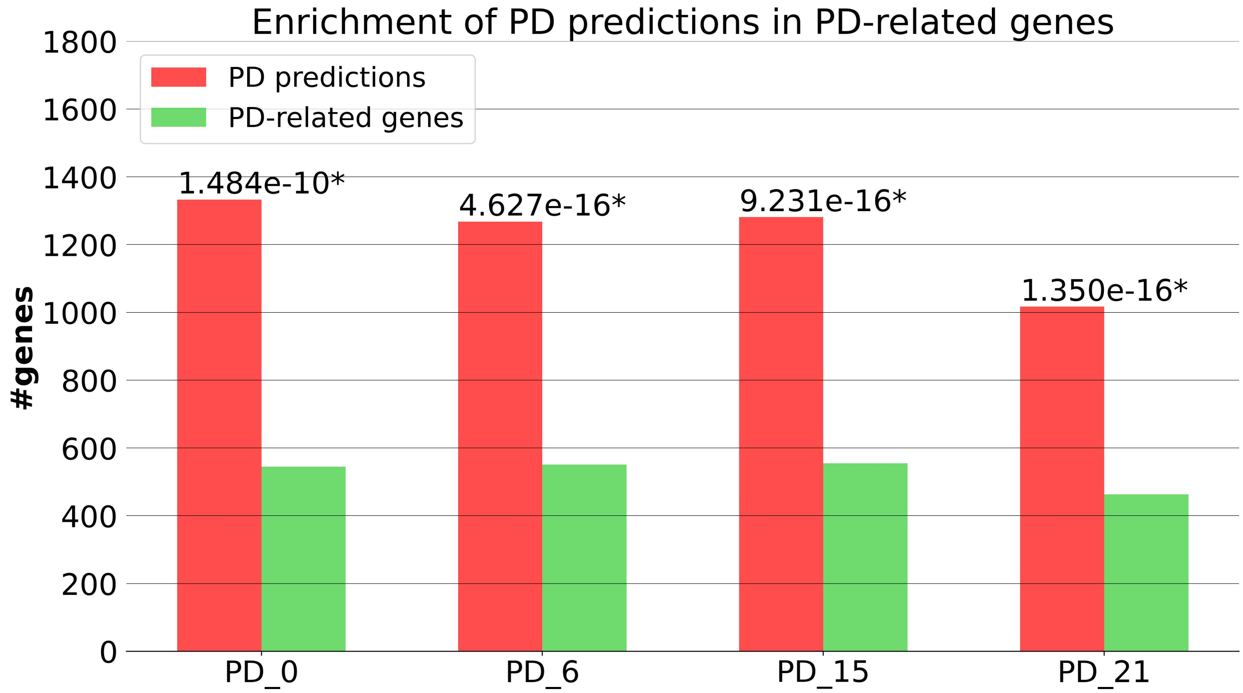


Supplementary Figure 10: **Enrichment of *Stage-specific PD predictions* in PD-related genes.** We perform the enrichment of ***Stage-specific PD predictions*** in PD-related genes (i.e., genes that co-occur with the term "Parkinson's disease" in at least one PubMed study, or are in the Gene4PD database). The bars represent the number of genes in a set of ***Stage-specific PD predictions*** (red) and the number of those genes that are in the PD-related gene set (green). The number above the bars for one set of ***Stage-specific PD predictions*** indicates the enrichment *p-value*. A set of predictions is enriched if *p-value* $< 0.05$, indicated by *.

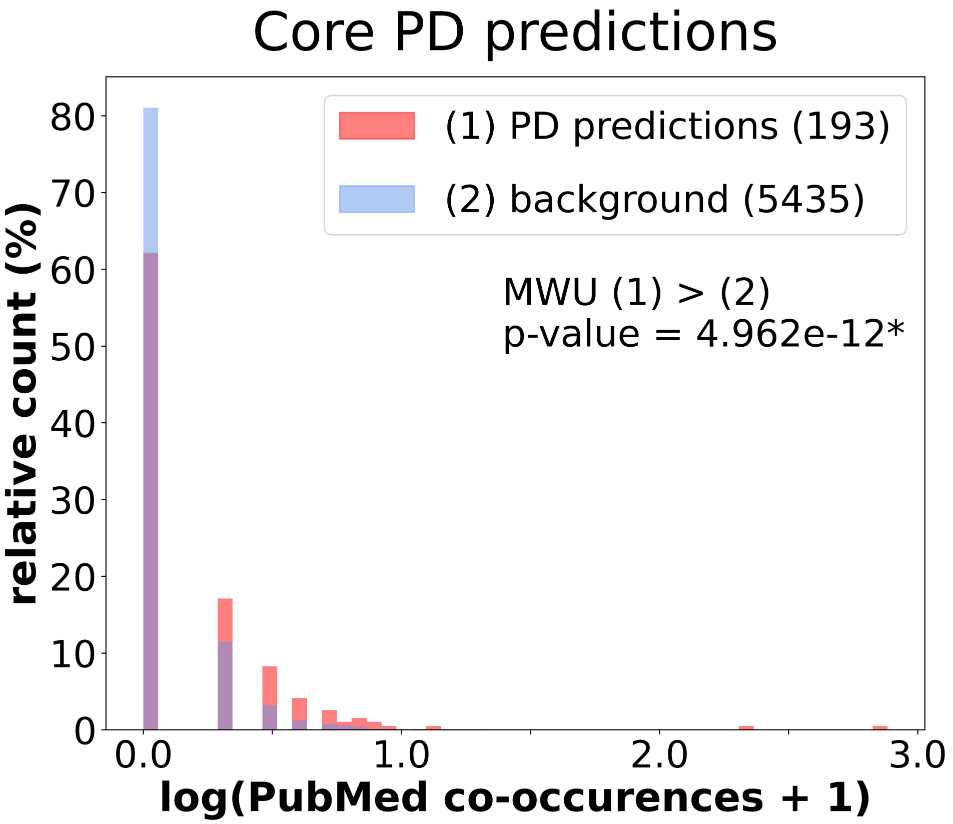


Supplementary Figure 11: **The log-transformed and normalized co-occurrence distributions of *Core PD predictions* with the term “Parkinson’s disease” in PubMed compared to the background set of genes.** A one-sided Mann-Whitney U (MWU) (with a significance level of 0.05) test checks if the distribution belonging to ***Core PD predictions*** is larger than for *background* (with *p-value* $< 0.05$ indicated by *). C: Control; PD: Parkinson’s disease; preds: predictions; num_PubMed: number of PubMed publications.

### Supplementary Tables

| CC | #genes | #SCs |
|---|---|---|
| $C_{D0}$ | 11,035 | 664 |
| $C_{D6}$ | 10,853 | 457 |
| $C_{D15}$ | 10,932 | 403 |
| $C_{D21}$ | 6,819 | 392 |
| $PD_{D0}$ | 9,263 | 507 |
| $PD_{D6}$ | 10,723 | 581 |
| $PD_{D15}$ | 10,654 | 392 |
| $PD_{D21}$ | 10,697 | 497 |

Supplementary Table 1: **Number of genes (#genes) and single cells (#SCs) of expression matrices for each cell condition (CC).** PD: Parkinson's disease; C: Control.

| CC | PPI | | COEX | | MI | | GI | |
|---|---|---|---|---|---|---|---|---|
| | #nodes | #edges | #nodes | #edges | #nodes | #edges | #nodes | #edges |
| $C_{D0}$ | 11,035 | 230,954 | 11,062 | 1,420,738 | 1,034 | 24,328 | 3,194 | 8,000 |
| $C_{D6}$ | 10,853 | 227,323 | 10,903 | 1,402,821 | 983 | 22,522 | 3,210 | 7,990 |
| $C_{D15}$ | 10,932 | 226,306 | 10,986 | 1,414,554 | 981 | 22,457 | 3,136 | 7,866 |
| $C_{D21}$ | 6,819 | 129,497 | 6,933 | 738,327 | 594 | 9,729 | 1,304 | 3,895 |
| $PD_{D0}$ | 9,263 | 193,843 | 9,311 | 1,157,162 | 866 | 17,820 | 2,796 | 7,230 |
| $PD_{D6}$ | 10,723 | 224,854 | 10,784 | 1,384,460 | 973 | 21,962 | 3,186 | 7,899 |
| $PD_{D15}$ | 10,654 | 220,123 | 10,718 | 1,375,439 | 960 | 21,892 | 3,092 | 7,749 |
| $PD_{D21}$ | 10,697 | 221,454 | 10,754 | 1,374,193 | 955 | 22,158 | 3,151 | 7,810 |

Supplementary Table 2: **Number of genes (#genes) and interactions (#edges) of each molecular network per cell condition (CC).** PD: Parkinson's disease; C: Control.

| CC | $k_1$ | $k_2$ | $mean_{\rho_{k_1},\rho_{k_2}}$ |
|---|---|---|---|
| $C_{D0}$ | 100 | 50 | 0.932 |
| $C_{D6}$ | 125 | 60 | 0.922 |
| $C_{D15}$ | 125 | 50 | 0.916 |
| $C_{D21}$ | 125 | 50 | 0.923 |
| $PD_{D0}$ | 75 | 60 | 0.905 |
| $PD_{D6}$ | 100 | 50 | 0.926 |
| $PD_{D15}$ | 100 | 60 | 0.911 |
| $PD_{D21}$ | 100 | 40 | 0.913 |

Supplementary Table 3: **Most optimal $k_1$, $k_2$ dimension parameters and mean dispersion coefficient ($mean_{\rho_{k_1},\rho_{k_2}}$) for each cell condition (CC).** PD: Parkinson's disease; C: Control.

| Rank | Gene | Evidence |
|---|---|---|
| 9 | PDIA6 | 35579911 |
| 19 | LMAN1 | 34930919 |
| 20 | GNAS | [16] |
| 28 | RPN2 | 16713278 |
| 44 | EGLN3 | 18069091 |
| 82 | PFKP | 35670764; Gene4PD |
| 144 | GOLT1B | 22542874 |
| 151 | FOS | 21507338 |

Supplementary Table 4: **Literature validation of the genes that are in the overlap between *Core PD predictions* and DEGs from the original study of the scRNA-seq data [17]**. The table ranks genes according to their average "movement" across all time points, so that genes with the largest "movement" are ranked at the top. The ID number in the evidence field is the PMID number of a study that shows why a prediction is relevant for PD. Genes also associated with PD in Gene4PD [13] are also annotated with Gene4PD in the Evidence field.

| Rank | Gene | Evidence | Selected Drug/Metabolite |
|---|---|---|---|
| 1 | PFN1 | 35628504 → 31493230 | Artenimol [30] |
| 2 | **CFL1** | 32819564 | |
| 3 | **DNAJC10** | 32662538; Gene4PD | |
| 4 | APLP2 | 34172567 → 31631455 | Zinc<br>Zinc-(acetate; chloride; sulfate) [30] |
| 5 | RRBP1 | GeneCards → 32854418 | Radezolid [30] |
| 6 | PTTG1IP | 34024830 → 28352155 | |
| 7 | **ENO1** | 15755676; Gene4PD | |
| 8 | RCN1 | 28319095 → 32854418 | Calcium [31] |
| 9 | **PDIA6** | 35579911; Gene4PD | |
| 10 | SEC63 | 34884562 → 32854418 | |
| 11 | CAPZB | GeneCards → 31493230 | |
| 12 | **TUBB** | 24275654; Gene4PD | |
| 13 | KDELR1 | GeneCards → 32854418 | |
| 14 | SOX4 | GeneCards → 32854418 | Progesterone [26] |
| 15 | SSR4 | GeneCards → 32854418 | Calcium [31] |
| 16 | ARPC2 | 20876399 → 31493230 | CK-636 [26] |
| 17 | **ALDOA** | 25626353; Gene4PD | |
| 18 | NREP | 28965931 | |
| 19 | **LMAN1** | 34930919 | |
| 20 | **GNAS** | [16] | |

Supplementary Table 5: **Validation of the top 20 *Core PD predictions* and their druggability**. The table ranks genes according to their average "movement" across all time points, so that genes with the largest "movement" are ranked at the top. The ID number in the evidence field is the PMID number of the study, showing why the prediction is relevant for PD. For studies where PMID is not available, we provide a citation. Genes in bold have literature that supports their role in PD, with some also associated with PD in Gene4PD [13] (annotated with Gene4PD in the Evidence field). For two PMIDs separated by an arrow in the Evidence field, the study labelled with the first PMID implicates the gene in a biological mechanism/function, and the second one explains how the mechanism/function is associated with PD. Genes whose basic function described in GeneCards [26] is PD-related have been annotated with GeneCards in the evidence field. The Selected Drug/Metabolite field provides compounds that can be used to target a particular gene.

| time point | Pubmed | | | PD gene enrichment | | |
|---|---|---|---|---|---|---|
| | 5% | 10% | StSp | 5% | 10% | StSp |
| D0 | $2.96e^{-07}$ | $2.73e^{-10}$ | $1.47e^{-09}$ | $1.33e^{-06}$ | $4.80e^{-07}$ | $3.19e^{-05}$ |
| D6 | $8.64e^{-12}$ | $7.40e^{-12}$ | $1.39e^{-11}$ | $2.41e^{-10}$ | $4.31e^{-09}$ | $3.50e^{-09}$ |
| D15 | $3.07e^{-10}$ | $5.65e^{-10}$ | $6.91e^{-09}$ | $6.73e^{-07}$ | $4.17e^{-08}$ | $3.96e^{-06}$ |
| D21 | $3.08e^{-03}$ | $1.75e^{-04}$ | $2.39e^{-04}$ | $2.71e^{-02}$ | $4.59e^{-03}$ | $7.49e^{-05}$ |
| Time points overlap | $3.83e^{-02}$ | $5.34e^{-03}$ | $6.99e^{-03}$ | $1.91e^{-02}$ | $3.20e^{-02}$ | $1.79e^{-02}$ |

Supplementary Table 6: **Literature validation of the predictions obtained by computing the "movement" between genes across time points and ranking them according to the largest "movement"**. The values in the table are p-values of two literature validation experiments checking if the predictions: 1) co-occur more with the term "Parkinson's disease" in Pubmed publications (column Pubmed), and 2) are significantly enriched in the PD-related genes (column PD gene enrichment). To obtain time point-specific predictions equivalent to the ***Stage-specific PD predictions*** (rows D0-D21) we threshold the ranked genes by taking the top 5% (column 5%) or 10% (column 10%) highest-ranked genes, or the same number of the highest-ranked genes as the ***Stage-specific PD prediction*** sets at matching time points (column StSp). Additionally, we perform the literature validation experiments on the individual overlaps of the sets of time-point specific predictions (row Time points overlap), representing ***Core PD predictions***.

#### References

[1] A. Anandhan, M. S. Jacome, S. Lei, P. Hernandez-Franco, A. Pappa, M. I. Panayiotidis, R. Powers, and R. Franco. Metabolic dysfunction in parkinson's disease: bioenergetics, redox homeostasis and central carbon metabolism. *Brain Research Bulletin*, 133:12–30, 2017.

[2] M. Ashburner, C. A. Ball, J. A. Blake, D. Botstein, H. Butler, J. M. Cherry, A. P. Davis, K. Dolinski, S. S. Dwight, J. T. Eppig, et al. Gene ontology: tool for the unification of biology. *Nature Genetics*, 25(1):25–29, 2000.

[3] Y. Benjamini and Y. Hochberg. Controlling the false discovery rate: a practical and powerful approach to multiple testing. *Journal of the Royal Statistical Society: series B (Methodological)*, 57(1):289–300, 1995.

[4] J.-P. Brunet, P. Tamayo, T. R. Golub, and J. P. Mesirov. Metagenes and molecular pattern discovery using matrix factorization. *Proceedings of the National Academy of Sciences*, 101 (12):4164–4169, 2004.

[5] C. H. Ding, T. Li, and M. I. Jordan. Convex and semi-nonnegative matrix factorizations. *IEEE Transactions on Pattern Analysis and Machine Intelligence*, 32(1):45–55, 2008.

[6] R. Ghemrawi and M. Khair. Endoplasmic reticulum stress and unfolded protein response in neurodegenerative diseases. *International Journal of Molecular Sciences*, 21(17):6127, 2020.

[7] V. Gligorijević, N. Malod-Dognin, and N. Pržulj. Patient-specific data fusion for cancer stratification and personalised treatment. In *Biocomputing 2016: Proceedings of the Pacific Symposium*, pages 321–332. World Scientific, 2016.

[8] A. Grover and J. Leskovec. node2vec: Scalable feature learning for networks. In *Proceedings of the 22nd ACM SIGKDD international conference on Knowledge discovery and data mining*, pages 855–864, 2016.

[9] B. Jassal, L. Matthews, G. Viteri, C. Gong, P. Lorente, A. Fabregat, K. Sidiropoulos, J. Cook,

M. Gillespie, R. Haw, et al. The reactome pathway knowledgebase. *Nucleic Acids Research*, 48(D1):D498–D503, 2020.

[10] M. Kanehisa, M. Furumichi, M. Tanabe, Y. Sato, and K. Morishima. Kegg: new perspectives on genomes, pathways, diseases and drugs. *Nucleic Acids Research*, 45(D1):D353–D361, 2017.

[11] H. Lee, W. S. James, and S. A. Cowley. Lrrk2 in peripheral and central nervous system innate immunity: its link to parkinson's disease. *Biochemical Society Transactions*, 45(1):131–139, 2017.

[12] L. Lestón Pinilla, A. Ugun-Klusek, S. Rutella, and L. A. De Girolamo. Hypoxia signaling in parkinson's disease: there is use in asking "what hif?". *Biology*, 10(8):723, 2021.

[13] B. Li, G. Zhao, Q. Zhou, Y. Xie, Z. Wang, Z. Fang, B. Lu, L. Qin, Y. Zhao, R. Zhang, et al. Gene4pd: A comprehensive genetic database of parkinson's disease. *Frontiers in Neuroscience*, 15, 2021.

[14] X. Ma, L. Liu, W. Yuan, Y. Zhang, and L. Song. Summary of static graph embedding algorithms. In *2023 4th International Conference on Computer Vision, Image and Deep Learning (CVIDL)*, pages 404–411. IEEE, 2023.

[15] N. Malod-Dognin, J. Petschnigg, S. F. Windels, J. Povh, H. Hemingway, R. Ketteler, and N. Pržulj. Towards a data-integrated cell. *Nature Communications*, 10(1):1–13, 2019.

[16] M. N. Murthy and N. B. Ramachandra. Prioritization of differentially expressed genes in substantia nigra transcriptomes of parkinson's disease reveals key protein interactions and pathways. *Meta Gene*, 14:12–18, 2017.

[17] G. Novak, D. Kyriakis, K. Grzyb, M. Bernini, S. Rodius, G. Dittmar, S. Finkbeiner, and A. Skupin. Single-cell transcriptomics of human ipsc differentiation dynamics reveal a core molecular network of parkinson's disease. *Communications Biology*, 5(1):1–19, 2022.

[18] F. Pedregosa, G. Varoquaux, A. Gramfort, V. Michel, B. Thirion, O. Grisel, M. Blondel,

P. Prettenhofer, R. Weiss, V. Dubourg, et al. Scikit-learn: Machine learning in python. *the Journal of Machine Learning Research*, 12:2825–2830, 2011.

[19] W. Poewe, K. Seppi, C. M. Tanner, G. M. Halliday, P. Brundin, J. Volkmann, A.-E. Schrag, and A. E. Lang. Parkinson disease. *Nature Reviews Disease Primers*, 3(1):1–21, 2017.

[20] N. Pržulj. *Analyzing Network Data in Biology and Medicine: An Interdisciplinary Textbook for Biological, Medical and Computational Scientists.* Cambridge University Press, 2019.

[21] H. Qiao. New svd based initialization strategy for non-negative matrix factorization. *Pattern Recognition Letters*, 63:71–77, 2015.

[22] R. Ray, J. K. Juranek, and V. Rai. Rage axis in neuroinflammation, neurodegeneration and its emerging role in the pathogenesis of amyotrophic lateral sclerosis. *Neuroscience & Biobehavioral Reviews*, 62:48–55, 2016.

[23] J. A. Rice. *Mathematical statistics and data analysis.* Cengage Learning, 2006.

[24] F. A. Scorza, A. C. Fiorini, C. A. Scorza, and J. Finsterer. Cardiac abnormalities in parkinson's disease and parkinsonism. *Journal of Clinical Neuroscience*, 53:1–5, 2018.

[25] R. J. Smeyne, A. J. Noyce, M. Byrne, R. Savica, and C. Marras. Infection and risk of parkinson's disease. *Journal of Parkinson's disease*, 11(1):31–43, 2021.

[26] G. Stelzer, N. Rosen, I. Plaschkes, S. Zimmerman, M. Twik, S. Fishilevich, T. I. Stein, R. Nudel, I. Lieder, Y. Mazor, et al. The genecards suite: from gene data mining to disease genome sequence analyses. *Current Protocols in Bioinformatics*, 54(1):1–30, 2016.

[27] T. Stuart, A. Butler, P. Hoffman, C. Hafemeister, E. Papalexi, W. M. Mauck III, Y. Hao, M. Stoeckius, P. Smibert, and R. Satija. Comprehensive integration of single-cell data. *Cell*, 177(7):1888–1902, 2019.

[28] `https://htmlpreview.github.io/?https://github.com/welch-lab/liger/blob/master/vignettes/Integrating_multi_scRNA_data.html`. Integrating multiple single-cell rna-seq datasets. accessed on June 2022.

[29] J. D. Welch, V. Kozareva, A. Ferreira, C. Vanderburg, C. Martin, and E. Z. Macosko. Single-cell multi-omic integration compares and contrasts features of brain cell identity. *Cell*, 177(7): 1873–1887, 2019.

[30] D. S. Wishart, Y. D. Feunang, A. C. Guo, E. J. Lo, A. Marcu, J. R. Grant, T. Sajed, D. Johnson, C. Li, Z. Sayeeda, et al. Drugbank 5.0: a major update to the drugbank database for 2018. *Nucleic Acids Research*, 46(D1):D1074–D1082, 2018.

[31] D. S. Wishart, A. Guo, E. Oler, F. Wang, A. Anjum, H. Peters, R. Dizon, Z. Sayeeda, S. Tian, B. L. Lee, et al. Hmdb 5.0: the human metabolome database for 2022. *Nucleic Acids Research*, 50(D1):D622–D631, 2022.
